# Asymmetric architecture and adaptation of *Treponema* flagella

**DOI:** 10.64898/2026.01.31.703064

**Authors:** Jiaqi Wang, Kurni Kurniyati, Wangbiao Guo, Jack M. Botting, Charles V. Sindelar, Chunhao Li, Jun Liu

**Affiliations:** Microbial Sciences Institute, Yale University, West Haven, CT, USA; Department of Microbial Pathogenesis, Microbial Sciences Institute, Yale University School of Medicine, New Haven, CT, USA; Philips Institute for Oral Health Research, School of Dentistry, Virginia Commonwealth University, Richmond, VA, USA; Department of Microbiology and Immunology, School of Medicine, Virginia Commonwealth University, Richmond, VA, USA; Yale Center for Research Computing, Yale University, New Haven, CT 06511

**Keywords:** Spirochetes, *Treponema*, Flagellin, Motility, Supercoiled bacterial motility

## Abstract

Spirochetes exhibit a distinctive corkscrew-like motility driven by periplasmic flagella that wrap around the cell body in a supercoiled configuration, yet the structural basis of this unique propulsion remains poorly understood. Here we combine cryo-electron microscopy, cryo-electron tomography, and genetic and biochemical analyses to determine the assembly and adaptation principles of the supercoiled flagellar filament in *Treponema denticola*, a major periodontal pathogen. Near-atomic structures reveal a glycosylated FlaB flagellin core encased by a previously unrecognized asymmetric sheath. The major sheath protein FlaA forms the bulk of the sheath and mechanically couples to the core through defined interfaces required for efficient motility, whereas four minor sheath proteins (FlaA1, FlaA2, FlaAP1, and FlaAP2) assemble along the concave side of the filament to accommodate intrinsic curvature. Disruption of this asymmetric core-sheath organization compromises force transmission and impairs motility, establishing coordinated asymmetric assembly as a fundamental mechanism underlying spirochetal motility.

## Main

Spirochetes are a distinct group of bacteria characterized by helical or flat-wave morphologies and a unique form of motility that enables efficient movement through highly viscous environments, including host tissues and mucus layers^1–3^. Several spirochetes are major human pathogens, including *Treponema denticola* (periodontitis), *Treponema pallidum* (syphilis), *Borreliella burgdorferi* (Lyme disease), *Leptospira* spp. (leptospirosis), and *Brachyspira* spp. (intestinal spirochetosis)^4–6^. Among these, the oral bacterium *Treponema denticola* is an obligatory anaerobic and motile pathogen implicated in both periodontal and endodontic infections and connects to several systemic illnesses including Alzheimer’s disease^7–9^. Motility, a central determinant of spirochetal pathogenicity, is driven by periplasmic flagella confined to the space between the outer membrane and peptidoglycan layer and wrapped around the cell body in a supercoiled form. Rotation of periplasmic flagella generates torsional forces that deform the cell body into helical or flat-wave conformations, producing a characteristic corkscrew-like motion that enables efficient migration through dense and heterogeneous environments^1,2^. In multiple species, genetic disruption of flagellar components abolishes infectivity, tissue penetration, and epithelial colonization, establishing motility as essential for host survival and transmission^10–15^. However, mechanisms underlying the unique spirochetal flagellar assembly and rotation have remained incompletely understood^1–3,16^.

The periplasmic flagellum is architecturally distinct from the external flagellum in model organisms such as *Escherichia coli* and *Salmonella enterica* although they all share a conserved organization consisting of a basal body, a hook, and a filament^16,17^. In spirochetes, however, the filament is uniquely composed of a conserved core encased by a distinct proteinaceous sheath^18^. The core is formed by flagellins homologous to *E. coli* and *S. enterica* FliC^19^ which are exported through the flagellar type III secretion systems (fT3SS)^18,20,21^. In contrast, the proteinaceous sheath is largely composed of FlaA proteins that are unique to spirochetes, lack homology to flagellins, and are exported via the Sec-dependent pathway rather than the fT3SS^18,22,23^.

Structural diversity among spirochetes is exemplified by *Leptospira*, whose flagellar filament contains an asymmetric, multi-protein sheath^24^. Disruption of genes involved in sheath formation not only impairs filament assembly but also compromises motility and infectivity^13,25,26^. Consistent with these observations, a recent high-resolution structure of mutant *Leptospira* flagellar filaments revealed pronounced asymmetry that is essential for proper filament assembly and motility^27^. Similarly, in *Brachyspira* and *Treponema* species, the filament consists of a core formed by three flagellins (FlaB1, FlaB2, and FlaB3) surrounded by a proteinaceous sheath primarily composed of FlaA^22,28–32^. Recent mass spectrometry analyses further demonstrated that *T. denticola* FlaB proteins are extensively O-glycosylated on serine and threonine residues, and that this modification is essential for filament assembly and motility^33^. Despite these advances, the structural basis and assembly mechanisms of sheathed flagella in spirochetes remain poorly understood.

Here, using cryo-electron microscopy (cryo-EM), we determine the near-atomic, asymmetric, multilayered architecture of the supercoiled flagellar filament in *T. denticola*. Our structure reveals that O-linked glycosylation of specific serine and threonine residues in FlaB flagellins stabilizes assembly of the filament core and promotes its interactions with the surrounding sheath. Structural analyses further show that the major sheath protein FlaA engages extensively with FlaB flagellins to form the bulk of the sheath, whereas four minor sheath proteins assemble selectively along the concave side of the supercoiled filament, accommodating the intrinsic curvature of the cell body. Finally, combining cryo-electron tomography (cryo-ET) with genetic and biochemical analyses, we demonstrate that deletion of any component of the asymmetric sheathed filament disrupts force transmission and severely impairs motility. Together, these findings establish a mechanistic framework for understanding how spirochetes have evolved an asymmetric periplasmic flagellar architecture to support their distinctive morphology and motility.

## Results

### Identification of novel flagellar proteins in *T. denticola*

Previous studies have shown that *T. denticola* periplasmic flagellar filaments are composed of three flagellins, including FlaB1 (TDE1477), FlaB2 (TDE1004), FlaB3 (TDE1475), and a major sheath protein, FlaA (TDE1712)^28,31–33^. Whole-cell cryo-ET revealed that two to three periplasmic flagella originate from each cell pole, wrap around the cell body in a supercoiled configuration, and extend toward the cell center, where filaments from opposite poles overlap (Fig. 1a)^32^. Notably, periplasmic flagella exhibit pronounced length asymmetry: filaments anchored at the old cell pole are longer than those at the new cell pole (Fig. 1a)^32^. Furthermore, filament diameter varies along its length, with the proximal region significantly thicker than distal regions, suggesting that sheath formation is not fully synchronized with filament core assembly (Fig. 1a,b)^32^. Consistent with this interpretation, deletion of *flaA* markedly reduces filament diameter along the entire cell length and severely impairs motility, underscoring the essential role of the sheath in flagellar architecture and function^32^.

**Fig. 1.**
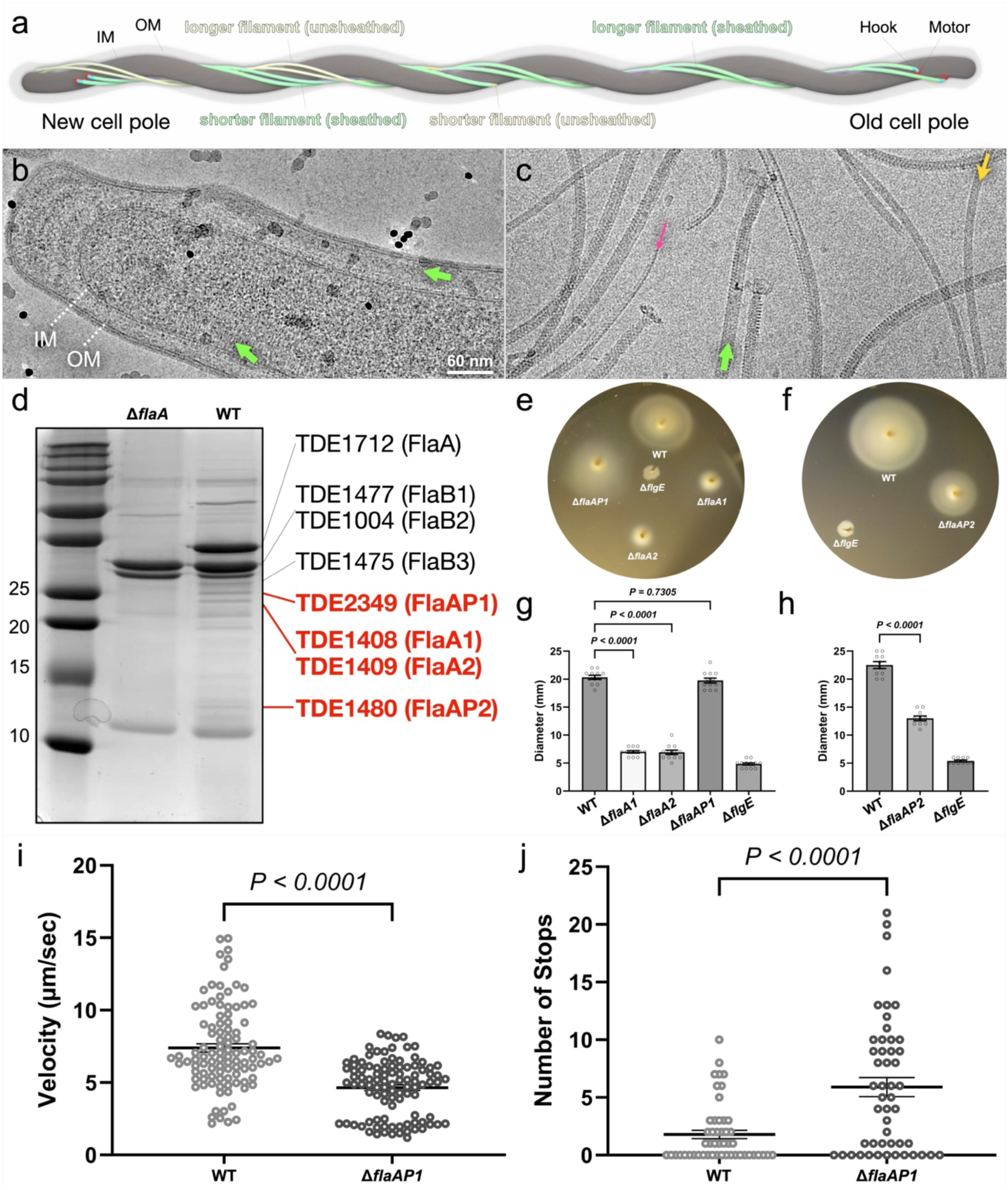
Identification and characterization of novel flagellar proteins in *T. denticola*. **a**, Schematic of *T. denticola* illustrating periplasmic flagella located between the inner and outer membranes, with variable filament lengths from the old and new cell poles. Notably, the sheath is colored in light green and the core is colored in yellow. Representative cryo-EM images from cell tips of wild-type (WT) cells (**b**) and purified flagellar filaments (**c**). **d,** Identification of new flagellar proteins of the *T. denticola* periplasmic flagella. SDS-PAGE analysis followed by Coomassie staining of purified periplasmic flagella from WT and Δ*flaA* strains. Bands indicated by red lines were excised and analyzed by nano LC-ESI/MS/MS for protein identification. The black-line-indicated bands represent periplasmic flagella-associated proteins that have been identified and studied. **e-h,** Swimming motility of WT, Δ*flgE*, Δ*flaA1*, Δ*flaA2,* Δ*flaAP1, and* Δ*flaAP2* cells in soft agar plates. Δ*flgE*, a non-motile mutant, was used as a control. **i,j,** Tracking analyses show a reduced velocity (**i**) and increased stops (**j**) of Δ*flaAP1* cells compared to WT cells.

To further define the protein composition of the flagellar sheath, periplasmic flagella were purified from wild-type and Δ*flaA* (Δ*TDE1712*) strains. Purified flagellar samples were examined by cryo-EM (Fig. 1c) and analyzed by SDS-PAGE followed by Coomassie staining (Fig. 1d). Cryo-EM imaging revealed heterogeneous filament populations with distinct diameters, including fully sheathed filaments, unsheathed filaments, and very thin protofilaments within the purified wild-type flagella (Fig. 1c).

SDS-PAGE analysis showed several protein bands present in wild-type samples but absent in the Δ*flaA* mutant (Fig. 1d, red lines). These bands were excised and subjected to nanoLC-ESI-MS/MS analysis, which identified two FlaA homologues, TDE1408 (236 aa, 27.0 kDa) and TDE1409 (246 aa, 27.8 kDa), hereafter designated FlaA1 and FlaA2, respectively. In addition, two previously uncharacterized FlaA-associated proteins were identified: TDE2349 (234 aa, 26.0 kDa) and TDE1480 (139 aa, 16.0 kDa), designated FlaAP1 and FlaAP2, respectively (Fig. 1d and Extended Data Fig. 1).

### Construction and functional characterization of new flagellar mutants

To understand the functions of these newly identified proteins, deletion mutants of *flaA1*, *flaA2*, *flaAP1*, and *flaAP2*, were generated by allelic exchange and further confirmed by PCR analysis (Extended Data Fig. 2). Immunoblotting revealed that levels of FlaA and FlaB proteins were not significantly altered in the Δ*flaA1* and Δ*flaA2* strains. In contrast, Δ*flaAP1* and Δ*flaAP2* strains exhibited a modest reduction in FlaA levels accompanied by a slight increase in FlaB abundance (Extended Data Fig. 3 and Extended Data Table 2,3).

Functional consequences of these deletions were assessed using swimming plate assays. The mean diameters of swimming rings (*n* = 6 plates) were 7.00 ± 0.21 mm for Δ*flaA1*, 6.92 ± 0.21 mm for Δ*flaA2*, and 13.00 ± 0.42 mm for Δ*flaAP2*, all significantly smaller than those of the wild-type strain (20.33 ± 0.36 mm to 22.50 ± 0.62 mm; P < 0.01) (Fig. 1e-h). We conclude that *flaA1*, *flaA2*, and *flaAP2* play important roles in *T. denticola* motility (Fig. 1e-h). In contrast, the mean diameter of swimming ring of Δ*flaAP1* was 19.75 ± 0.45 mm slightly smaller than that in wild type (Fig. 1e,g and Extended Data Table 3). To better compare the motility phenotypes of wild-type and Δ*flaAP1* cells, we deployed motion-tracking analysis to show that the Δ*flaAP1* cells have reduced swimming velocity compared to wild-type cells in 1% methylcellulose (Fig. 1i; Movie S1,2). In addition, the Δ*flaAP1* cells have more stops compared to wild-type cells (Fig. 1j; Movie S1,2). Together, these data indicate that FlaAP1 also plays a role in *T. denticola* motility.

### Near-atomic asymmetric structures of the supercoiled flagellar filament in *T. denticola*

To define the molecular architecture of the supercoiled flagellar filament, we used single-particle cryo-EM to analyze filaments both *in situ* and *in vitro* (Extended Data Fig. 4-7). Asymmetric reconstruction and refinement in cryoSPARC^34^ yielded near-atomic structures of the supercoiled filament (Fig. 2a-d and Extended Data Fig. 4-7). The filament core consists of 11 flagellin protofilaments (C1-C11) with an overall diameter of 125 Å and a central lumen diameter of 24.5 Å. The core structure is similar to a filament with a helical symmetry of a 4.80 Å rise and a 65.29° twist (Fig. 2d, g). However, each flagellin protofilament is slightly different from the adjacent ones to allow formation of the supercoiled filament (Fig. 2 and Extended Data Fig. 7). The filament sheath exhibits pronounced asymmetric features with an overall diameter of ∼195 Å (Fig. 2a, d). Specifically, the sheath is mostly formed by eight protofilaments (S1-S8) of the major sheath protein FlaA and six protofilaments (AS1-AS6) of four minor sheath proteins (Fig. 2h). Consistent with flagellar proteins identified by nanoLC-ESI-MS/MS analysis (Fig. 1), the near-atomic map enabled *de novo* model building of the filament core protein FlaB1, the major sheath protein FlaA, four minor sheath proteins (FlaA1, FlaA2, FlaAP1, and FlaAP2) (Fig. 2e-h and Extended Data Fig. 6a-d). The overall model reveals a molecularly heterogeneous flagellar architecture, providing a structural framework for understanding the specialized adaptations of the supercoiled periplasmic flagella and their contributions to the distinctive morphology and motility of spirochetes (Fig. 2i-m and Extended Data Fig. 8).

**Fig. 2.**
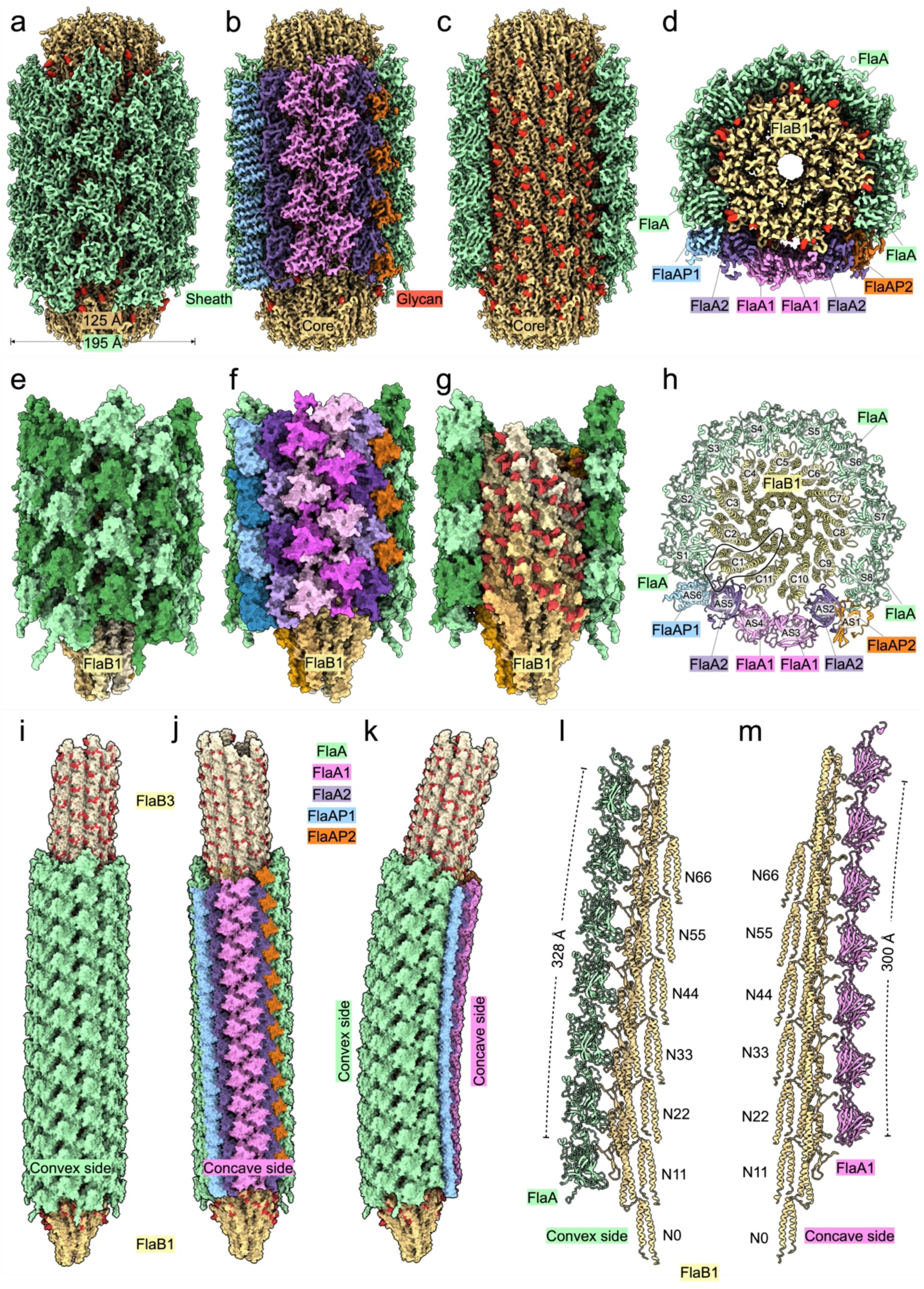
Asymmetric architecture of the supercoiled flagellar filament in *T. denticola*. **a,b**, Cryo-EM structure of the sheathed flagellar filament viewed from the symmetric (**a**) and asymmetric (**b**) sides. **c**, Close-up view of the glycosylated filament core. Filament components are colored by protein identity; glycans are shown in red. **d**, Perpendicular cross-section of the filament revealing a central lumen, a tubular filament core, and a surrounding protein sheath. The core is predominantly formed by FlaB1, whereas the sheath is composed of five proteins: FlaA, FlaA1, FlaA2, FlaAP1, and FlaAP2. **e-h**, Asymmetric model of the sheathed filament shown from the symmetric side (**e**), asymmetric side (**f**), and the glycosylated filament core (**g**), with a perpendicular view of a single-layer filament model shown in (**h**). **i-k,** Three different views of the architecture of the supercoiled periplasmic flagellar filament formed by three core flagellins (FlaB1-FlaB3), five sheath proteins (FlaA, FlaA1, FlaA2, FlaAP1, and FlaAP2). **l,** The models of the FlaB1 protofilament and FlaA protofilament at the convex side. The total length of the 7-subunit FlaA protofilament is 328Å. **m,** The models of the FlaB1 protofilament and FlaA2 protofilament at the concave side. The total length of the 7-subunit FlaA1 protofilament is 300Å. The FlaA1 protofilament is slightly shorter than the FlaA protofilament.

### The core of the sheathed filament is mainly formed by glycosylated FlaB1

Assembly of the flagellar filament core in *T. denticola* is mediated by three FlaB flagellins that share highly similar sequences and overall structures but differ in their stoichiometries in the wild-type and in various *flaB* mutants^32^. For example, deletion of *flaB3* increases the ratio of FlaB1 to FlaA from 0.71:1 to 1.12:1, while reducing the ratio of FlaB2 to FlaA from 0.61:1 to 0.36:1^32^. These observations suggest that FlaB1 is the major flagellin and is critical for assembly of the sheathed flagellar filament core.

Consistent with this hypothesis, our cryo-EM analysis of purified sheathed flagella reveals that the core of the sheathed filament is composed of 11 FlaB1 protofilaments (Fig. 3). Each FlaB1 subunit consists of the conserved D0 and D1 domains (Fig. 3a). Moreover, in agreement with previous mass spectrometry analyses^33^, five serine or threonine residues (Ser115, Ser126, Thr137, Ser152, and Thr171) are modified by glycans corresponding to 2-methoxy-4,5,6-trihydroxy-hexanoyl-L-glycero-L-manno-nonulosonic acid (Fig. 3a-c and Extended Data Fig. 6e). These glycan modifications cluster around three surface-exposed loops in the D1 domain (loops 1-3), where they appear to stabilize inter-subunit interactions within the filament core (Fig. 3d,e). In addition, FlaB1 protofilaments assemble into a supercoiled filament through conserved longitudinal interactions mediated by residues D106, E92, R91, Q88, and E225, as well as lateral interactions involving residues R223, R244, and N277 (Fig. 3f, g).

**Fig. 3.**
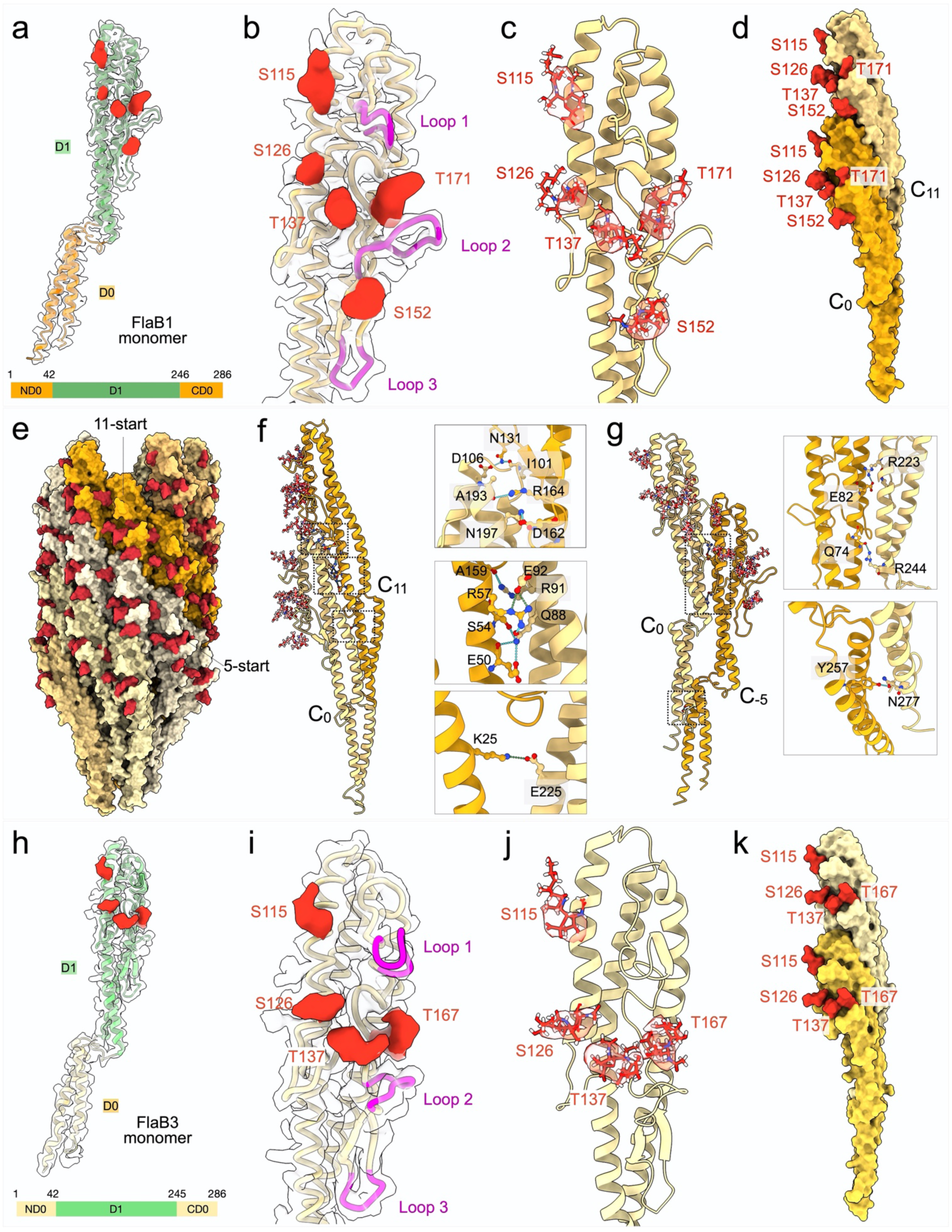
The flagellar filament core is composed of glycosylated FlaB proteins assembled into a conserved helical structure. **a**, Atomic model of FlaB1 fitted into a single subunit of the cryo-EM map of the sheathed filament core at 2.87 Å resolution. Five extra densities corresponding to glycans are colored in red. **b,** Five O-glycosylation sites identified in the D1 domain of FlaB1, distributed around three flexible surface loops (magenta). **c**, Refined models of glycans at residues S115, S126, T137, S152, and T171 fitted in the corresponding glycan densities. **d**, Longitudinal FlaB1 dimer with glycans shown in red. The glycan attached to S152 appears involved in the interaction between two FlaB1 molecules. **e**, Overall architecture of the glycosylated FlaB1 filament core. **f,** Overview of the longitudinal interactions between two FlaB1 molecules along the 11-start protofilament (left panel). Detailed side-chain interactions (right panel). **g**, Overview of the lateral interactions between two FlaB1 molecules along the 5-start protofilament (left panel). Detailed side-chain interactions (right panel). **h**, Model of FlaB3 fitted into the single subunit map from the unsheathed filament core at 2.87 Å resolution. Four extra densities corresponding to glycans are colored in red. **i**, Four O-glycosylation sites identified in FlaB3 D1domain, distributed around three flexible surface loops (magenta). **j**, Refined model of the FlaB3 glycans at residues S115, S126, T137, and T167 fitted in the corresponding glycan densities. **k**, Longitudinal FlaB3 dimer with glycans shown in red. Notably, there is no glycan for S152 in FlaB3.

Given the essential role of glycosylation in flagellar filament assembly and motility in *T. denticola*^33^, these findings elucidate how specific glycan modifications and conserved inter-subunit interactions cooperate to drive filament core assembly and maintain the mechanical stability required to accommodate elastic deformation and supercoiling.

### Non-sheathed filaments in wild-type cells are formed by glycosylated FlaB3

Comparative analysis of *in situ* structures of sheathed and non-sheathed filaments in wild-type *T. denticola* cells reveals that the non-sheathed filament is predominantly composed of glycosylated FlaB3. Similar to FlaB1, FlaB3 contains the D0 and D1 domains, which mediate formation of the filament core through conserved inter-subunit interactions (Fig. 3h).

Consistent with mass spectrometry analyses^33^, FlaB3 is glycosylated at four residues: Ser115, Ser126, Thr137, and Thr167 (Fig. 3i, j). Each glycan is clearly resolved in the cryo-EM maps (Fig. 3i-k), These glycans are positioned on surface-exposed regions of the filament core, suggesting a role in stabilizing inter-subunit contacts.

Because non-sheathed filaments are consistently observed at the distal portion of flagella in wild-type *T. denticola* cells, these findings suggest that the three flagellins are sequentially assembled during flagellar biogenesis, with FlaB1 and FlaB2 incorporated at early stages and FlaB3 added later.

### The major sheath protein FlaA is intimately coupled to the FlaB1 core

Our cryo-EM map enabled construction of an atomic model of the major sheath protein FlaA, revealing how it assembles into the bulk of the filament sheath through extensive longitudinal and lateral interactions (Fig. 4a, b). Longitudinal contacts are mediated by a combination of electrostatic and hydrophobic interactions involving D293, D220, and D271 from one FlaA subunit and R104, R292, and R306 from an adjacent subunit (Fig. 4c). Lateral interactions are formed by salt bridges involving an extended peptide region, with key residues K109, K280, and R239 interacting with E345, E342, and E328 on neighboring subunits (Fig. 4d).

**Fig. 4.**
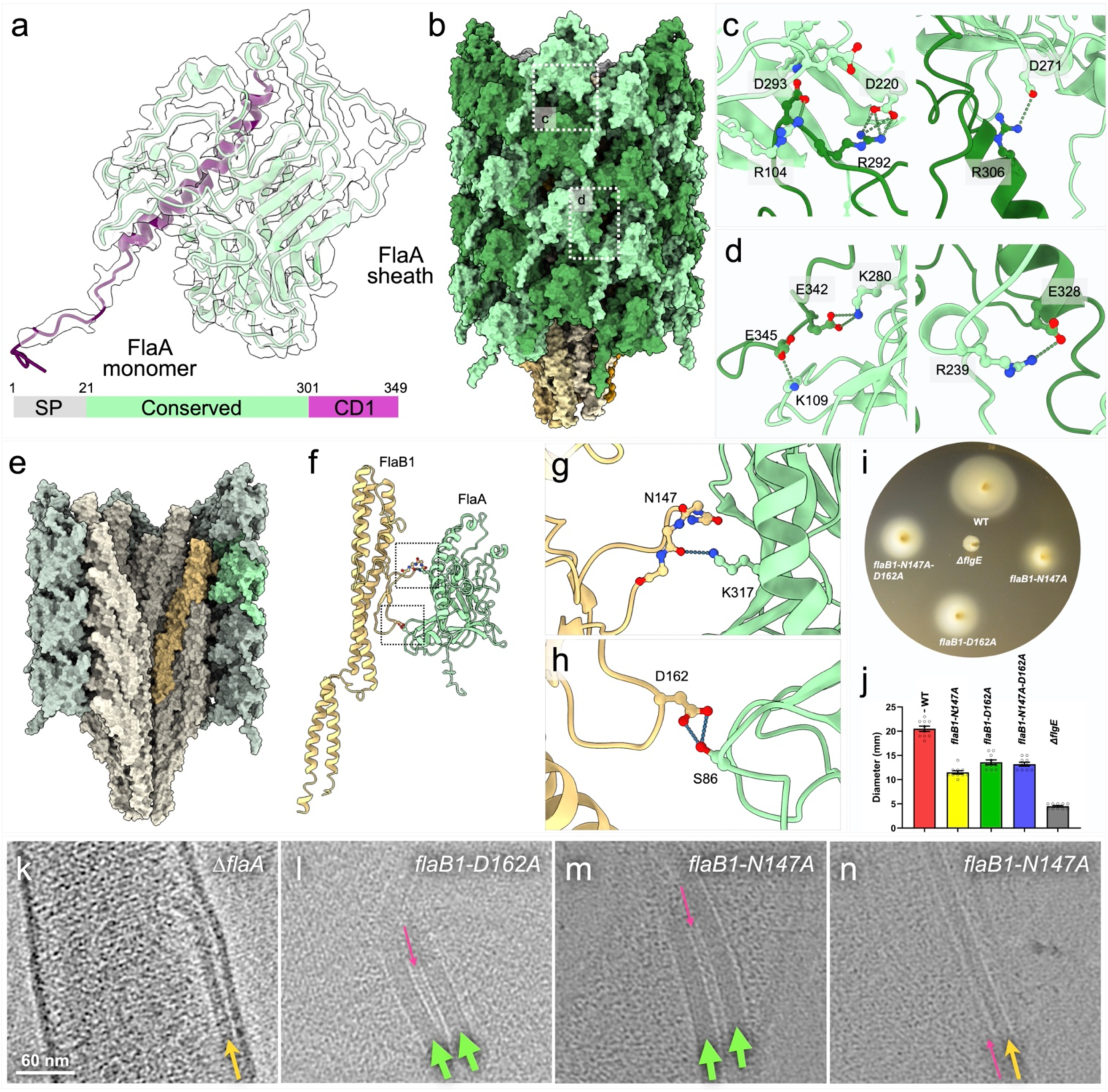
The *T. denticola* flagellar sheath is predominantly composed of FlaA proteins. **a,** Atomic model of FlaA fitted into a single FlaA subunit of the cryo-EM map. **b**, Overview of the filament sheath formed by FlaA. **c,d,** Detailed views of longitudinal (**c**) and lateral (**d**) interactions between adjacent FlaA subunits. **e**, Cross-sectional view of the sheathed filament model. One FlaA and one FlaB1 monomer are highlighted in brighter colors. **f,** The FlaA-FlaB1 model showing interactions between FlaA and FlaB1. **g,h,** Detailed views of key interactions: Asn147 of FlaB1 interacting with Lys317 of FlaA (**g**), and Asp162 of FlaB1 interacting with Ser86 of FlaA (**h**). **i,j**, Swimming plate assays showing that site-directed mutations in *flaB1* significantly impair cell motility. Swimming ring diameters (*n* = 6 plates) for *flaB1-N147A* (11.50 ± 0.34 mm), *flaB1-D162A* (13.60 ± 0.50 mm), and flaB1-N147A/D162A (13.20 ± 0.39 mm) are significantly smaller (P < 0.01) than those of wild-type cells (20.50 ± 0.54 mm). **k-n**, Cryo-ET reconstructions reveals filament with variable diameters in Δ*flaA* (**k**), *flaB1-D162A* (**l**), and *flaB*-N147A mutants (**m,n**). Sheathed filaments (Thick) are labeled with green arrows, unsheathed filaments are labeled with yellow arrows, and very thin protofilaments are labeled with purple arrows.

Importantly, each FlaA subunit directly engages FlaB1 to form a repeating core-sheath composite unit that propagates along the filament core (Fig. 4e). At the FlaB1-FlaA interface, specific interactions were observed between N147 of FlaB1 and K317 of FlaA, as well as between D162 of FlaB1 and S86 of FlaA (Fig. 4f-h), indicating an intimate mechanical coupling between the FlaB1 core and the FlaA sheath.

### FlaB1 site-directed mutants destabilize the filament sheath and impair motility

To test whether the two identified FlaB1-FlaA interactions are required for flagellar assembly and spirochetal motility, residues N147 and D162 of FlaB1 were individually or combinatorially substituted with alanine. Mutations were introduced by in-frame replacement of the wild-type *flaB1* gene with the corresponding mutant alleles, as schematized in Extended Data Fig. 2e (top). The resulting strains were validated by PCR (Extended Data Fig. 2e, bottom) and DNA sequencing. The three point mutants were designated *flaB1*-N147A, *flaB1*-D162A, and *flaB1*-N147A-D162A, respectively.

Immunoblot analysis showed that levels of both FlaA and FlaB proteins were largely unchanged in the *flaB1*-N147A-D162A double mutant, slightly reduced in the *flaB1*-N147A mutant, and modestly increased in the *flaB1*-D162A mutant (Extended Data Fig. 2e and Extended Data Table 2,3). Despite these modest differences in protein abundance, all three mutants exhibited pronounced motility defects relative to the wild-type strain (Fig. 4i, j and Extended Data Table 3). Swimming plate assays revealed significantly reduced swimming ring diameters (*n* = 6 plates) for *flaB1*-N147A (11.50 ± 0.34 mm), *flaB1*-D162A (13.60 ± 0.50 mm), and *flaB1*-N147A-D162A (13.20 ± 0.39 mm), compared with wild type (20.50 ± 0.54 mm; *P* < 0.01) (Fig. 4i, j).

To further assess the structural consequences of these mutations, cryo-ET was used to compare *flaB1*-D162A and *flaB1*-N147A cells with the Δ*flaA* mutant. As expected, Δ*flaA* filaments were thin and unsheathed (Fig. 4k)^32^. In contrast, filaments in both *flaB1*-D162A and *flaB1*-N147A cells were predominantly thick and sheathed (Fig. 4l-n). Notably, thin protofilaments were also observed in these mutants, indicating that although FlaA is able to assemble into a sheath, individual sheath protofilaments can partially detach from the filament core (Fig. 4k-n and Extended Data Table 3).

Together, these results demonstrate that FlaB1 residues N147 and D162 are critical for robust interactions between the flagellar core and sheath. Although these core-sheath interactions are not strictly required for sheathed filament assembly, they are essential for maintaining core-sheath stability, thereby enhancing the mechanical integrity of the sheathed filament and promoting efficient motility in *T. denticola*.

### FlaA1 and FlaA2 assemble along the concave side of the supercoiled filament

The concave side of the supercoiled filament is associated with two minor sheath proteins: FlaA1 and FlaA2 (Fig. 5a). Both FlaA1 and FlaA2 are homologs of FlaA (Fig. 5b,c and Extended Data Fig. 1). Notably, two FlaA1 protofilaments assemble into a central, “Tai-chi”-like longitudinal dimer, flanked bilaterally by two FlaA2 protofilaments (Fig. 5a). Stabilization of this asymmetric extension is mediated by FlaA1-FlaA1 interactions involving residues E197, K203, and D221 from one subunit and R176, D227, and H72 from the adjacent subunit (Fig. 5d), while FlaA2 provides additional longitudinal reinforcement (Fig. 5e-g).

**Fig. 5.**
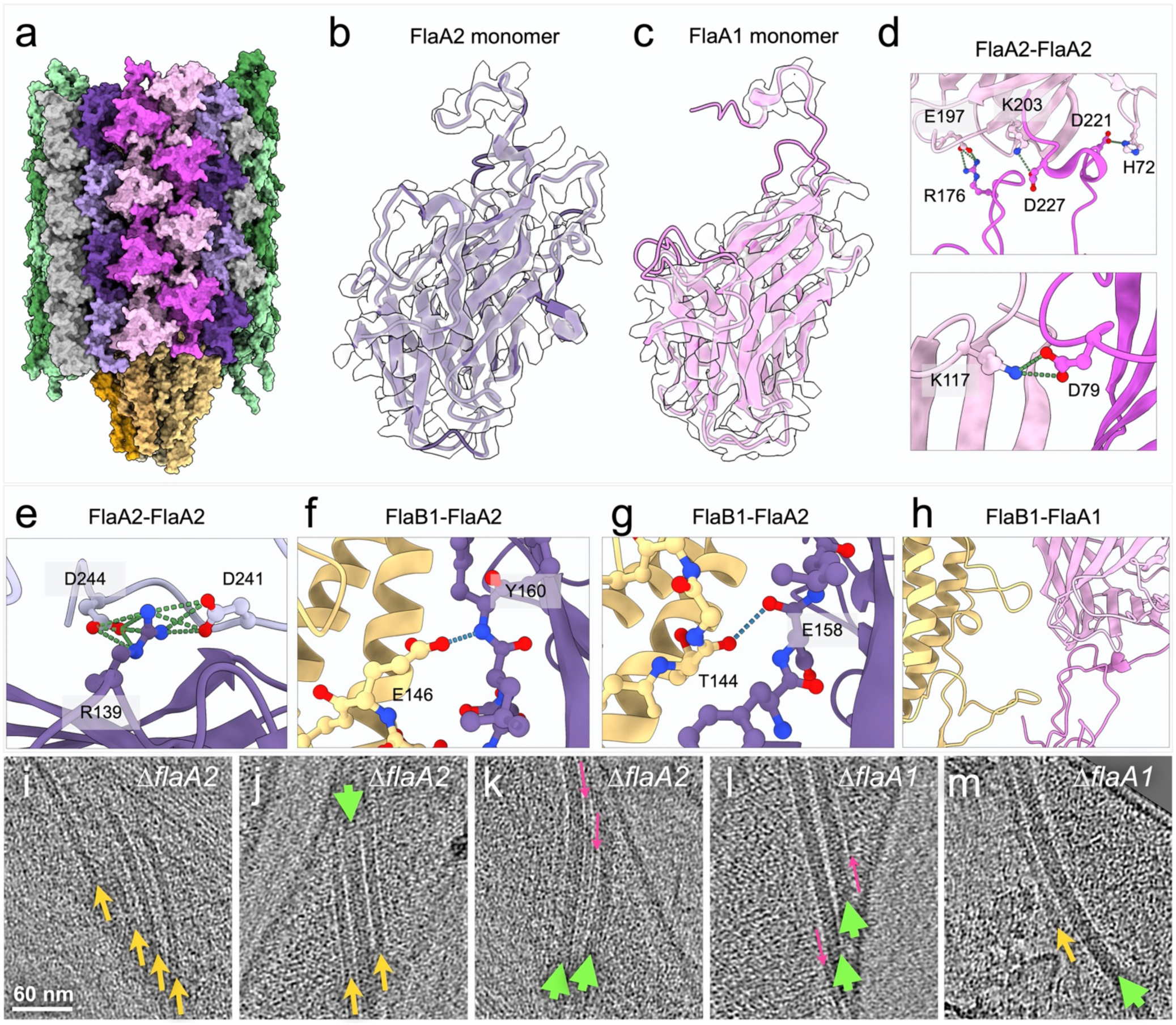
FlaA1 and FlaA2 form the concave side of the supercoiled filament in *T. denticola*. **a**, Surface view of the concave side of the supercoiled filament sheath, mainly composed of FlaA1 and FlaA2. **b,c**, Atomic models of FlaA2 (**b**) and FlaA1 (**c**) fitted into their corresponding cryo-EM maps of the filament sheath. **d**, Detailed views of FlaA1 dimer interactions, showing longitudinal (top) and lateral (bottom) contacts. **e**, Detailed view of longitudinal dimer interactions between FlaA2 subunits. **f,g,** Close-up views of key interactions between FlaA2 and FlaB1, including Glu146 of FlaB1 interacting with Tyr160 of FlaA2 (**f**) and Thr144 of FlaB1 interacting with Glu158 of FlaA2 (**g**). **h**, No specific interaction between FlaB1 and FlaA1 is observed. **i-k**, Cryo-ET reconstructions of the Δ*flaA2* mutant reveal filaments with variable diameters, indicating the presence of sheathed filaments (green arrows), unsheathed filaments (yellow arrows), and protofilaments (purple arrows). **l,m**, Cryo-ET reconstructions of the Δ*flaA1* mutant similarly show filaments with variable diameters, indicating the coexistence of sheathed filaments (green arrows), unsheathed filaments (yellow arrows), and protofilaments (purple arrows).

In contrast to the strong FlaA-FlaB1 interactions, interactions between FlaA2 and FlaB1 are markedly weaker, and no direct interaction between FlaA1 and FlaB1 is detected (Fig. 5f-h), indicating reduced core-sheath coupling along the concave side of the supercoiled sheathed filament. Cryo-ET analyses of Δ*flaA1* and Δ*flaA2* mutants reveal filaments with variable diameters, consistent with the ability of both mutants to assemble sheathed filaments but with reduced sheath stability. This instability results in the frequent presence of unsheathed filaments and detached protofilaments. Importantly, both mutant strains exhibit pronounced motility defects (Fig. 1e-h), demonstrating that FlaA1 and FlaA2 are critical contributors to the mechanical integrity and structural stability of the supercoiled, sheathed flagellar filament.

### FlaA associated proteins FlaAP1 and FlaAP2 strengthen the supercoiled filament

FlaAP1 and FlaAP2 are two previously unrecognized sheath-associated proteins that localize to the concave side of the sheathed filament (Fig. 5a). FlaAP1 is a HEAT-repeat protein^35^ that assembles into a longitudinal chain stabilized by extensive hydrophobic interactions and specific hydrogen bonds between N208 and K196 of one subunit and E53 and Y37 of the adjacent subunit (Fig. 6a,b). Occupation of this position by FlaAP1 induces a rearrangement of the terminal peptide of FlaA2 (Fig. 6c,d).

**Fig. 6.**
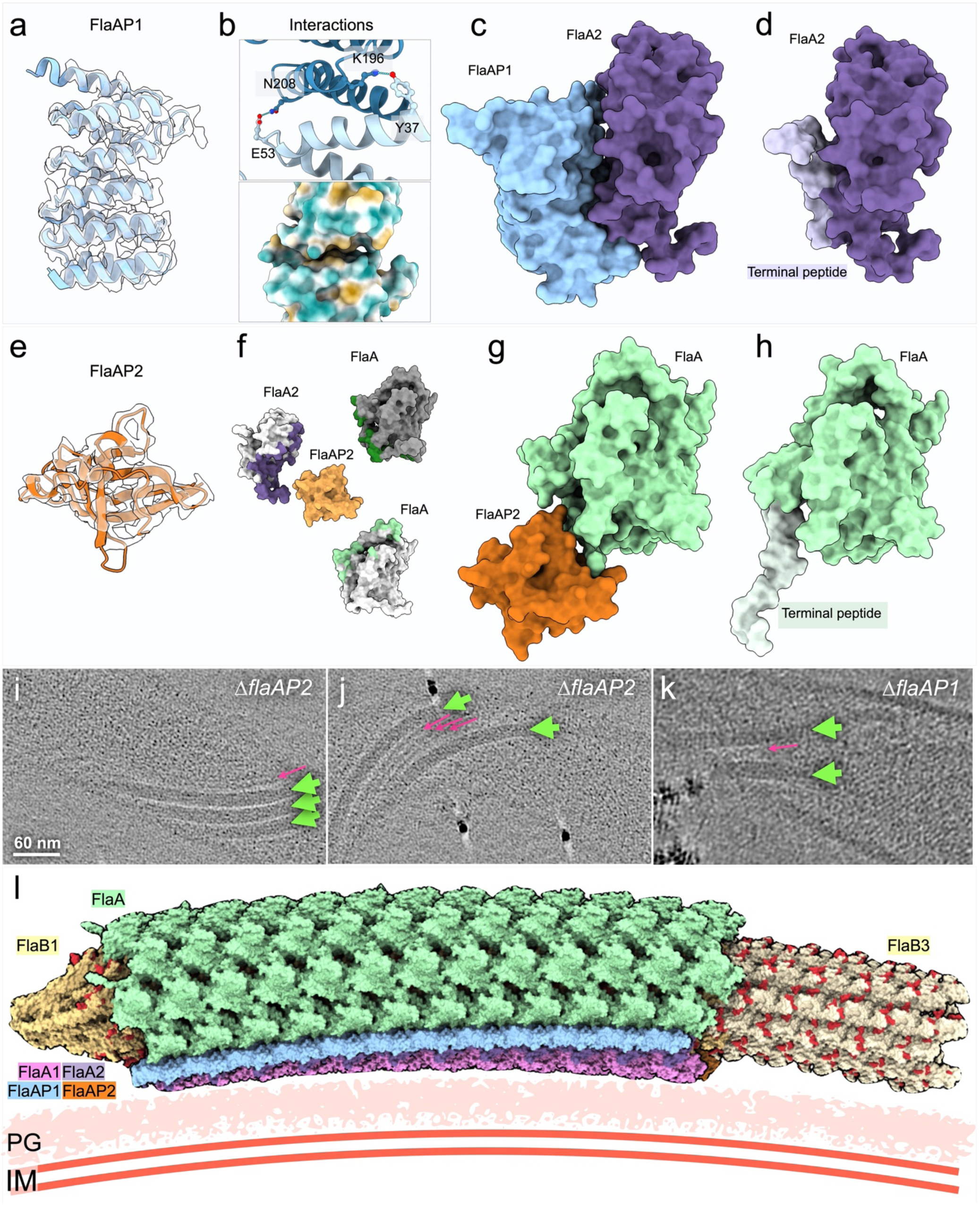
FlaA-associated proteins FlaAP1 and FlaAP2 assemble along the concave side of the supercoiled filament. **a**, Atomic model of FlaAP1 fitted into a single FlaAP1 subunit map. **b,** Interfaces of two adjacent FlaAP1 molecules. Top, longitudinal contacts between adjacent subunits; bottom, hydrophobic interactions stabilizing the FlaAP1 dimer. **c,d**, FlaAP1 binding induces a conformational rearrangement in the adjacent FlaA2. A flexible peptide segment (lighter color), which occupies a lateral position in the absence of FlaAP1 (**d**), shifts outward to accommodate FlaAP1 binding (**c**). **e,** Atomic model of FlaAP2 fitted into a single FlaAP2 subunit map. **f,** Interactions of FlaAP2 with three neighboring subunits, including FlaA2, longitudinal FlaA, and lateral FlaA. **g,h**, FlaAP2 binding alters the lateral FlaA conformation. A flexible peptide segment (lighter color) relocates from its original position (**h**) to enable FlaAP2 binding (**g**). **i-k**, Cryo-ET reconstructions reveals filaments with variable diameters in Δ*flaAP2* (**i,j**) and Δ*flaAP1* (**k**) mutant cells. **l,** Model illustrating the assembly of four minor sheath proteins along the concave side of the supercoiled filament while the major sheath protein FlaA forms the rest of the sheath, enabling the filament to wrap tightly around the cell body. PG: Peptidoglycan, IM: Inner membrane.

FlaAP2 adopts a distinctive “diamond”-like fold and interacts simultaneously with three neighboring subunits—FlaA2, a longitudinal FlaA, and a lateral FlaA (Fig. 6e,f). Notably, interaction between FlaAP2 and the longitudinal FlaA induces a rearrangement of the terminal peptide of FlaA (Fig. 6g,h). In both cases, the corresponding terminal peptide densities of FlaA and FlaA2 are not resolved in the presence of FlaAP1 or FlaAP2, likely reflecting increased conformational flexibility.

Cryo-ET analyses reveal that sheathed flagellar filaments are present in both Δ*flaAP1* and Δ*flaAP2* mutants; however, numerous thin protofilaments are frequently observed detached from the filament core, indicating reduced sheath stability in both mutants (Fig. 6i–k). Consistent with these structural defects, both mutant strains exhibit significantly impaired motility (Fig. 1e–j), implicating FlaAP1 and FlaAP2 as key contributors to the structural stability and mechanical performance of the supercoiled flagellar filament in *T. denticola*.

Taken together, longitudinal cross-sectional views of the sheathed filament highlight extensive hydrophobic regions at interlayer interfaces, forming hydrophobic contacts that contribute to overall filament integrity (Extended Data Fig. 9). Representative hydrophobic interactions within FlaA longitudinal dimers and within FlaAP1 longitudinal dimers further underscore the importance of hydrophobic packing in maintaining filament architecture (Extended Data Fig. 9). Electrostatic surface analyses of the sheathed filament reveal distinct charge distributions across the structure (Extended Data Fig. 10). The outer surface of the sheath is predominantly negatively charged, whereas positively charged regions on the inner surface of the sheath are positioned adjacent to the negatively charged, glycosylated outer surface of the filament core (Extended Data Fig. 10). This complementary charge distribution is consistent with electrostatic contributions to core-sheath association and stabilization of the sheathed filament.

## Discussion

To infect and disseminate within mammalian hosts, spirochetes have evolved a morphology and mode of motility uniquely adapted for propulsion through viscous media and dense tissue barriers^1^. Central to this adaptation are periplasmic flagella, which reside within the periplasmic space and are fundamentally distinct from the external flagella. Rather than extending into the extracellular milieu, spirochetal flagella wrap tightly around the cell body in an intrinsically supercoiled configuration, and their rotation drives coordinated deformation and propulsion of the entire cell, enabling efficient motility in environments where external flagella are ineffective^16^. Here, through an integrated structural, genetic, and biochemical approach, we define the asymmetric and multilayered architecture of the supercoiled flagellar filament in *T. denticola*. The filament core is composed of multiple glycosylated flagellins with distinct spatial distributions, encased within an asymmetric sheath in which the major sheath protein FlaA forms most of the surface, while four minor sheath proteins assemble preferentially along the concave side of the supercoil (Fig. 6l; Movie S3). Importantly, genetic disruption of any sheath component compromises motility (Fig. 1e-h), demonstrating that this architectural asymmetry is not merely structural but functionally essential.

### How does the asymmetric, multilayered flagellar filament assemble?

The presence of extended non-sheathed distal regions supports a model in which flagellar core assembly precedes sheath formation. Flagellins are exported by the fT3SS^18,20,21^ and polymerize at the distal tip of the nascent filament to form a heterogeneous, supercoiled core, with distinct flagellin isotypes occupying spatially segregated regions along the filament axis. Similar organization has been recently observed in the sheathed flagella of *Vibrio cholerae*^36^ and *Helicobacter pylori*^37^, suggesting that differential flagellin incorporation is a common strategy during filament assembly. However, unlike the membranous sheaths of *V. cholerae* and *H. pylori*, which are derived from the outer membrane, the periplasmic flagellar core is encased by sheathed proteins that are independently exported via the Sec secretion pathway^18,22,23^ and polymerize onto the pre-formed supercoiled core (Movie S4). Glycosylation and flexible surface-exposed loops on the core facilitate intimate core-sheath interactions that likely enhance filament rigidity and torsional resistance.

### How does rotation of the asymmetric, multilayered flagellar filament drive spirochetal motility?

The asymmetric, multilayered architecture of the periplasmic flagellar filament has direct consequences for spirochetal hydrodynamics by generation of propulsive forces^38,39^. Such asymmetry stabilizes the supercoiled geometry during rotation and promotes efficient transmission of motor-generated torque. Rather than functioning as an isolated propeller, rotation of the periplasmic filament drives deformation and counter-rotation of the entire cell body. Coordinated rotation of periplasmic flagella anchored at opposite poles converts motor torque into whole-cell corkscrew motion (Movie S4), transforming viscous resistance into productive thrust and providing a mechanistic basis for efficient spirochetal motility in highly viscous host environments.

### How are the assembly and rotation of the flagellar filament regulated?

The mechanical performance of the periplasmic flagellar filament is tightly coupled to its genetic regulation. In *T. denticola*, four flagellar filament genes are partitioned between two transcriptional programs controlled by σ⁷⁰ and σ²⁸, providing temporal and quantitative control over the production of distinct filament components^32,40^. This layered regulation likely governs the sequence and stoichiometry of filament assembly, thereby fine-tuning filament length, thickness, stiffness, and core-sheath coupling. Because these morphological parameters are directly linked to distinct swimming mechanisms, as demonstrated in spirochetes such as *Treponema*, *Leptospira*, and *Borrelia*^3,16^, regulatory control over filament composition directly translates into control of rotational dynamics, torque transmission, and swimming behavior.

In sum, our studies reveal a uniquely asymmetric architecture of the supercoiled periplasmic flagellar filament that is required for tight wrapping around the cell body, efficient torque transmission, and whole-cell deformation. We establish the periplasmic flagellar filament as a mechanically integrated and genetically tunable organelle optimized for rotation-driven propulsion, providing a structural and regulatory framework for understanding spirochetal motility in complex environments.

## Methods

### Bacterial strains, culture conditions, and oligonucleotide primers

*T. denticola* ATCC 35405 (wild type, WT) was used in this study^31^. Bacterial cells were grown in either tryptone-yeast extract-gelatin-volatile fatty acids-serum (TYGVS) medium or Oral Bacterial Growth Medium (OBGM) at 37°C in an anaerobic chamber in the presence of 90% nitrogen, 5% carbon dioxide, and 5% hydrogen^41,42^. *T. denticola* isogenic mutants were grown with appropriate antibiotic for selective pressure as needed: erythromycin (50 µg/ml) or gentamicin (20 µg/ml). *Escherichia coli* 5α strain (New England Biolabs, Ipswich, MA) was used for DNA cloning. The *E. coli* strains were cultivated in lysogeny broth (LB) supplemented with appropriate concentrations of antibiotics for selective pressure as needed: ampicillin (100 µg/ml). The oligonucleotide primers for PCR amplifications used in this study are listed in Extended Data Table 1. These primers were synthesized by IDT (Integrated DNA Technologies, Coralville, IA).

### Periplasmic flagella isolation

Isolation of the periplasmic flagella was performed as previously described ^32^. Briefly, 500 ml of the mid-logarithmic (log)-phase *T. denticola* cultures (∼5×10^8^ cells/ ml) were centrifuged at 5,000×*g* for 20 min at 4°C. The cells were washed four times with phosphate-buffered saline (PBS, pH 7.4) and once with T1 buffer (0.15 M Tris-HCl buffer, pH 6.8). The final pellets were resuspended in 30 ml of T1 buffer. Three milliliters of 10% Triton X-100 were slowly added, mixed and incubated for 1 hour at room temperature. And then, 3 ml of 200 µg/ml mutanolysin (Sigma-Aldrich, St. Louis, MO) was added slowly, followed by addition of 300 µl of T2 buffer (0.1 M Tris-HCl buffer, pH 6.8). The resultant mixture was first incubated for 2 hours at room temperature and then at 4°C overnight. After the incubation, 600 µl of 0.1 M MgSO4 was added, followed by addition of 600 µl of T2 buffer. The mixture was incubated for 5 min at room temperature and then centrifuged at 17,000×*g* for 15 min at 4°C. The supernatant containing periplasmic flagella was collected and 2 ml of 20% PEG 8000 (Alfa Aesar, London, UK) in 1 M NaCl was added and then incubated for 30 min on ice. The resultant sample was centrifuged at 27,000×*g* for 30 min at 4°C. The pellet containing periplasmic flagella was resuspended in alkaline solution (0.1 M KCl, 0.5 M sucrose, 0.1% Triton X-100, 50 mM sodium bicarbonate, pH 11) and incubated for 1 hour on ice. The periplasmic flagella were finally collected by centrifugation at 80,000×*g* for 45 min at 4°C and washed once in 20 mM Tris-HCl buffer, pH 8. The flagellar pellet was resuspended in water and stored at 4°C for further analyses.

### Electrophoresis and immunoblotting analyses

Sodium dodecyl sulfate-polyacrylamide gel electrophoresis (SDS-PAGE) and immunoblotting analyses were carried out as previously described ^43^. For immunoblotting analysis, *T. denticola* cells were harvested in the mid-log phase (∼5×10^8^ cells/ml) Equal amounts of whole-cell lysates (∼5 μg) were separated on SDS-PAGE. The resultant gels were subjected to Coomassie blue staining or immunoblotting analyses. The antibodies against *T. pallidum* FlaBs and *T. denticola* FlaA and DnaK are previously described^33,44^. The stoichiometry of each flagellar filament protein was analyzed using Image Lab software from Bio-Rad (Bio-Rad).

### Constructions of *T. denticola* flagellar mutants

The diagrams in Extended Data Fig. 2 illustrate the deletion of the open reading frames (ORFs) of *TDE1408*, *TDE1409*, *TDE2349*, and *TDE1480*. *TDE1408* is used as an example to describe how this vector (*TDE1408::ermB*) was constructed and used to delete the target gene. To construct *TDE1408::ermB*, the *TDE1408* upstream flanking region (UR) and a previously described erythromycin B resistant cassette (*ermB*)^45^ were PCR amplified with primers P1/P2 and P3/P4, respectively, and then fused together with primers P1/P4 to generate UR-ermB. The downstream flanking region of *TDE1408* (DR) was PCR amplified with primers P5/P6, and subsequently fused to UR-ermB fragment by PCR using primers P1/P6. The final fragment, UR-ermB-DR, was cloned into the pGEM-T easy vector (Promega, Madison, WI), generating *TDE1408∷ermB* (Extended Data Fig. 2a). The same method was used to construct *TDE1409::ermB* and *TDE2349::ermB*.

To construct plasmid (*TDE2349::ERM*) to delete *TDE2349*, the following method was used. The *TDE2349* upstream flanking region (UR) and the downstream flanking region (DR) were PCR amplified with primers P13/P14 and P17/P18, respectively, and then fused together with primers P13/P18 to generate UR-DR fragment. The UR-DR fragment was cloned into the pMD19 T-vector (Takara Bio USA, Inc, Mountain View, CA). The full-length erythromycin cassette was PCR amplified with primers P15/P16, generating Fragment ERM, which was subsequently cloned into the pGEM-T easy vector (Promega). The Fragment UR-DR and ERM were digested using NotI and then ligated, generating the *TDE2349::ERM* plasmid. The resulting constructs were transformed into *T. denticola* wild-type competent cells via electroporation and then plated on soft agar containing erythromycin, as previously described ^40^. The deletions were confirmed by PCR (Extended Data Fig. 2, bottom panels). The resultant mutants were designated as Δ*TDE1408*, Δ*TDE1409*, Δ*TDE2349*, and Δ*TDE1480*, respectively.

### Construction of *flaB1* site-directed mutants in *T. denticola*

The vector (*FlaB1::ermB*) shown in Extended Data Fig. 2e was constructed to replace the wild-type *flaB1* (*TDE1477*) with genes carrying site-directed mutations at amino acids N147A, D162A, or both N147A and D162A, using two-step PCR and DNA cloning as previously described ^40^. Briefly, to construct this plasmid, the upstream flanking region of FlaB1 along with the wild-type *flaB1* gene (UR), *ermB*, and the downstream flanking region of *flaB1* (DR) were PCR amplified using primers P28/P29, P30/P31, and P32/P33, respectively. Then, UR and *ermB* were PCR ligated using P28/P31 to generate the UR-*ermB* fragment, which was subsequently PCR ligated to DR using primers P28/P33. The final PCR product, UR-ermB-DR, was cloned into the pJET1.2 vector (Thermo Fisher Scientific, San Jose, CA). Site-directed mutagenesis of *flaB1* in this plasmid was performed using the Q5 Site-Directed Mutagenesis Kit (New England Biolabs) according to the manufacturer’s instructions. The primers used to replace N147 and N162 with Ala were P35/P36 and P37/P38, respectively. The resulting mutations were confirmed by DNA sequencing analysis. The resulting constructs were transformed into *T. denticola* wild-type competent cells via electroporation and then plated on soft agar containing erythromycin, as previously described^40^. Site-directed mutations in the resulting colonies were confirmed by PCR followed by DNA sequencing.

### Bacterial swimming plate assay and motion tracking analysis

A swimming plate assay of *T. denticola* was performed as previously described^32^. Swimming plate assays were performed with plates containing 50% (v/v) TYGVS:50% (v/v) PBS with 0.35% (w/v) SeaPlaque agarose. Briefly, 10 ml of the late-log phase *T. denticola* cultures (10^9^ cells/ ml) were harvested by centrifugation at 5,000×g for 10 min at room temperature. The cell pellets were resuspended in ∼40 µL of spent culture supernatant. Each plate was inoculated with 4 µL of resuspended cell cultures and incubated anaerobically at 37°C for 2-3 days to allow the cells to swim out. The plates were incubated anaerobically at 37°C for 3-5 days to allow the cells to swim out. The diameters of the swimming rings were measured in millimeters. As a negative control, a previously constructed *T. denticola* non-motile mutant, Δ*flgE*, was included to determine the initial inoculum size^46^. The average diameters of each individual strains were calculated from three or six independent plates; and the results are represented as the mean of diameters ± standard error of the mean (SEM).

For motion-tracking analysis, 100 μl of mid-log phase *T. denticola* cultures was first diluted (1:1) in OBGM medium and then 10 μl of diluted cultures was mixed with an equal volume of 2% methylcellulose with a viscosity of 4,000 cp (MC4000). *T. denticola* cells were videotaped and edited using a computer-based bacterial tracking system alongside the software Velocity (Improvision Inc., Coventry, UK), as previously described^32^. For each bacterial strain, the cells were recorded for up to 30 sec.

### In-gel trypsin digestion

The excised bands were cut into ∼2 mm cubes and subjected to in-gel digestion 47,48. The excised gel pieces were washed consecutively with 200 µL deionized water, followed by 200 µL 50mM ammonium bicarbonate in water/50% acetonitrile (ACN) and finally 200 µL 100% ACN. The dehydrated gel pieces were dried in a speed vacuum (SpeedVac SC110 Thermo Savant, Milford, MA) and reduced with 10mM DTT (dithiothreitol) (w/v) in 100mM ammonium bicarbonate in water for 1 hour at 60°C, then alkylated by adding 55mM iodoacetamide in 100mM ammonium bicarbonate (w/v) and incubation at room temperature, in the dark, for 45 minutes. Wash steps were repeated as described above. The gel pieces were dried in a speed vacuum and rehydrated with 100 µl trypsin (Promega Sequencing Grade) at 15ng/µl (w/v) in 50mM ammonium bicarbonate/10% ACN on ice for 20 minutes, topped with 150µl 50mM ammonium bicarbonate, 10% ACN (just until gel was covered), and incubated at 37°C for 16 hours. The digestion was stopped by addition of 10 µl 100% formic acid (FA) in water, incubated at room temperature for 5 minutes and the supernatant transferred to a clean 1.5ml polypropylene low-bind microfuge tube. The gel pieces were then extracted twice by adding 150μl of 50% ACN/5% FA and vortexing at 1800rpm for 30 minutes followed by sonication for 5 minutes, and once by adding 150μl of 75% ACN/5% FA with incubation at room temperature for 5 minutes. The extraction supernatants were pooled with the first supernatant, dried in a speed vacuum. The final sample was reconstituted into 2% ACN/0.5% FA and filtered through a 0.22 µm cellulose acetate spin filter (Cornin75Costar Spin-X) prior to nanoLC-MS/MS analysis.

### Protein Identification by nano LC/MS/MS Analysis

The analysis was carried out using an Orbitrap Fusion^TM^ Tribrid^TM^ (Thermo Fisher Scientific) mass spectrometer equipped with a nanospray Flex Ion Source, and coupled with a Dionex UltiMate 3000 RSLCnano system (Thermo, Sunnyvale, CA) ^47^. The peptide samples (10 μL) were injected onto a PepMap C-18 RP viper trapping column (5 µm, 100 µm i.d x 20 mm) at 20 µL/min flow rate for rapid sample loading and then separated on a PepMap C-18 RP nano column (2 µm, 75 µm x 25 cm) at 35 °C. The tryptic peptides were eluted in a 90-min gradient of 5% to 35% ACN in 0.1% formic acid at 300 nL/min, followed by an 8-min ramping to 90% ACN-0.1% FA and an 8-min hold at 90% ACN-0.1% FA. The column was re-equilibrated with 0.1% FA for 25 min prior to the next run. The Orbitrap Fusion was operated in positive ion mode with spray voltage set at 1.2 kV and source temperature at 275°C. External calibration for FT, IT and quadrupole mass analyzers were performed. In data-dependent acquisition (DDA) analysis, the instrument was operated using FT mass analyzer in MS scan to select precursor ions followed by 3 second “Top Speed” data-dependent CID ion trap MS/MS scans at 1.6 m/z quadrupole isolation for precursor peptides with multiple charged ions above a threshold ion count of 10,000 and normalized collision energy of 30%. MS survey scans at a resolving power of 120,000 (fwhm at m/z 200), for the mass range of m/z 300-1600. Dynamic exclusion parameters were set at 50 s of exclusion duration with ±10 ppm exclusion mass width. All data were acquired under Xcalibur 4.3 operation software (Thermo-Fisher Scientific).

### Data analysis

The DDA raw files with MS and MS/MS were subjected to database searches using Proteome Discoverer (PD) 2.4 software (Thermo Fisher Scientific) with the Sequest HT algorithm. The PD 2.4 processing workflow containing an additional node of Minora Feature Detector for precursor ion-based quantification was used for protein identification and relative quantitation of identified peptides. The database search was conducted against *T. denticola* ATCC 35405 NCBI database that contains 3704 sequences. The peptide precursor tolerance was set to 10 ppm and fragment ion tolerance was set to 0.6 Da. Oxidation of M, deamidation of N and Q were specified as dynamic modifications of amino acid residues; protein N-terminal acetylation, M-loss and M-loss plus acetylation were set as variable modifications, carbamidomethyl C was specified as a static modification. Only high confidence peptides defined by Sequest HT with a 1% FDR by Percolator were considered for confident peptide identification.

### Cryo-EM sample preparation

For *in situ* data, *T. denticola* cells were centrifuged at 2,000g for ∼5 minutes in 1.5 mL tubes, and the resulting pellet was gently rinsed with phosphate-buffered saline (PBS). The cell pellet was resuspended in PBS to a final optical density (OD600) of 1.0 for plunge-freezing preparation. For *in vitro* data, 4 µl aliquot of purified flagellar filaments was applied onto a freshly glow-discharged grid (200 mesh, R2/1, Quantifoil). The grids were blotted with filter paper (Whatman^TM^) for ∼6s and rapidly plunged into a liquid ethane-propane mixture using a GP2 plunger (Leica). During the plunge-freezing process, the GP2 environmental chamber was maintained at 25°C and 90% relative humidity to optimize vitrification.

### *In-situ* cryo-EM data collection and processing

Cryo-EM specimen was imaged using a 300 kV Titan Krios electron microscope (Thermo Fisher Scientific) equipped with a field emission gun and a post-GIF K3 Summit direct electron detector (Gatan). Images were recorded in nanoprobe mode with the following settings: magnification at 81,000×, spot size at 5, illumination aperture at 1.22 µm, slit width at 20 eV, C2 aperture at 50 or 70 µm, and objective aperture at 100 µm. A customized multishot script was used to directly image the cell tips on the cryo-EM grids with SerialEM^49^, applying defocus values from −1.6 µm to −2.2 µm. The total electron dose was ∼70 e^−^/Å^2^. The effective pixel size was 1.068 Å per physical pixel (super-resolution mode, half physical pixel size). A total of 6,672 micrographs were collected from *T. denticola* wild-type cells.

All micrographs were first motion-corrected^50^ and then subjected to Patch CTF Estimation^51^ in cryoSPARC^34^. Filament particles were automatically picked using the Filament Tracer^52^ in cryoSPARC with a filament diameter of 400 Å, a minimum inter-particle distance of 0.15 Å, and a minimum filament length of 1 Å. Selected particles were extracted with a box size of 720 pixels and Fourier-binned to 360 pixels. 2D classification was used to remove non-filament and low-quality particles, yielding a dataset of 379,837 particles.

An initial filament model was generated by *ab initio* reconstruction, followed by heterogeneous and local refinements for 3D reconstruction. Helical symmetry parameters, including helical rise and twist, were determined using the Symmetry Search Utility in cryoSPARC^34^. The top-ranked solution, with a helical rise of 4.80 Å and a twist of 65.29°, was selected. Using these parameters, the final helical reconstruction of the sheathed flagellar filament in *T. denticola* reached a near-atomic resolution of 3.12 Å.

Subsequently, 3D classification was performed using a sheath-focused mask. Three distinct classes were identified based on sheath features: symmetric sheath, asymmetric sheath, and non-sheathed filaments. Particles from each class were subjected to local refinement to further improve resolution. During refinement, the rotational search range was progressively reduced from 20° to 2°, the translational search ranges from 10 Å to 2 Å, and the initial low-pass filter from 12 Å to 3 Å. This process yielded three final maps corresponding to symmetric sheath, asymmetric sheath, and non-sheathed filaments, containing 256,356 (after particle expansion), 206,104, and 123,481 particles, with FSC (0.143) resolutions of 3.12 Å, 3.51 Å, and 2.36 Å, respectively.

To investigate the spatial distribution of the sheath along filament curvature and within the cell body, well-aligned particles were re-extracted using a larger box size of 1,280 pixels and Fourier-binned to 448 pixels. 2D classification was then performed, and classes exhibiting pronounced curvature and double-membrane features, corresponding to the inner membrane of the cell body, were selected for Homogeneous Reconstruction only. The resulting low-resolution 3D map was visualized in ChimeraX^53^, revealing that the asymmetric sheath preferentially localizes to the inner curvature of the filament and appears oriented toward the inner membrane. In parallel, selected particles were mapped back onto the original micrographs by Manual Picker to confirm their spatial distribution in the raw data.

All cryo-EM maps were further processed using the deep learning-based refinement algorithm DeepEMhancer^54^ to enhance visualization.

### *In-vitro* cryo-EM data collection and processing

Cryo-EM specimen of purified flagellar filaments from *a T. denticola* Δ*flaB3* mutant was imaged using a 300 kV Titan Krios electron microscope as described above. The total electron dose was ∼50 e^−^/Å^2^. A total of 4319 micrographs were collected from purified flagellar filaments of the *T. denticola* Δ*flaB3*.

All micrographs were first motion-corrected^50^ and then subjected to Patch CTF Estimation^51^ in cryoSPARC. Filament particles were automatically picked using the Filament Tracer^52^ in cryoSPARC with a filament diameter of 240 Å, a minimum inter-particle distance of 0.1 Å, and a minimum filament length of 2 Å. Selected particles were extracted with a box size of 360 pixels. 2D classification was used to remove non-filament and low-quality particles, yielding a dataset of 643,096 particles.

Homogeneous and local refinement was used for 3D reconstruction, using a ∼20 Å lowpass-filtered *in situ* cryo-EM map as the initial model. Subsequently, 3D classification was performed using a sheath-focused mask. Three distinct classes were identified based on sheath features: disrupted or low-quality sheath, asymmetric sheath, and non-sheathed filaments. 328,259 particles in disrupted or low-quality sheath class were excluded for the reconstruction. 181,566 particles from asymmetric sheath were further classified using sheath-focused mask by 3D classification until each class displayed significant asymmetric features. Maps from each class were aligned using the Align 3D Maps in cryoSPARC^34^, with particle alignments updated to place the asymmetric region at a consistent location. All the well-aligned particles were refined by homogeneous refinement and followed by local refinements for 3D reconstruction. 133,271 particles from non-sheathed class were subjected to local refinement to further improve resolution.

During local refinements for all the classes, the rotational search range was progressively reduced from 20° to 2°, and the translational search ranges from 10 Å to 2 Å. This process yielded two final maps corresponding to asymmetric sheath and non-sheathed filaments, containing 181,566, and 133,271 particles, with FSC (0.143) resolutions of 3.12 Å, and 3.47 Å, respectively.

### Cryo-ET data collection and analysis

Frozen-hydrated specimens of *T. denticola* cells were imaged on a 200kV Glacios transmission electron microscope (Thermo Fisher Scientific). Tilt series were acquired using SerialEM^49^ in conjunction with the FastTomo script^55^ and PaceTomo algorism^56^, employing a dose-symmetric acquisition scheme. Imaging was conducted at a defocus of −4.8 µm, with a cumulative electron dose of ∼70 e^−^/Å^2^ distributed evenly across 33 tilts, ranging from −48° to +48° in 3° increments. Raw frames were corrected for beam-induced motion using MotionCorr2^50^, followed by tilt alignment and stacking in IMOD^57^. Tomographic reconstructions were generated from 4× binned tilt series using Tomo3D^58^. In total, 71 tilt series were acquired for seven mutants (**Extended Data Table 4**).

### Model building

The amino acid sequences of FlaB1, FlaB2, FlaB3, FlaA, FlaA1, FlaA2, FlaAP1, and FlaAP2 were retrieved from UniProt (Accession numbers, FlaB1: Q73MN1; FlaB2: Q73NZ6; FlaB3: Q73MN3; FlaA: Q73M00; FlaA1: Q73MU9; FlaA2: Q73MU8; FlaAP1: Q73K73; FlaAP2: Q73MM8) and used as input for AlphaFold3^59^ to generate predicted structural models. The predicted models were fitted into the cryo-EM map using ChimeraX^53^, followed by interactive manual adjustment in Coot^60^ and real-space refinement in Phenix^61^. The final monomer models exhibited excellent stereochemistry, as evaluated by PHENIX. Phenix reported a map-to-model correlation coefficient of 0.70 for Δ*flaB3* sheathed filament, 0.72 for Δ*flaB3* non-sheathed filament, 0.68 for WT non-sheathed filament, which served as robust cross-validation metrics for evaluating map quality and the accuracy of C1 symmetry parameters. Furthermore, several residues extended beyond the cryo-EM density, indicating poor model-to-map fit and the local flexibility. Sequence alignments were performed using Clustal Omega^62^. ChimeraX^53^ was used for cryo-EM map segmentation, visualization, and molecular surface rendering. Interface and interaction analyses among residues were performed using PDBe PISA^63^. ChimeraX^53^ and Blender (www.blender.org) were used for making animations.

## Data availability

The atomic coordinates and corresponding density maps of the flagellins have been deposited in the Protein Data Bank (PDB) and the Electron Microscopy Data Bank (EMDB). The sheathed filament from the *T. denticola* ATCC 35405 Δ*flaB3* strain is available under PDB: 10PL and EMD-75374. The non-sheathed filament from the same strain is deposited as PDB: 10PM and EMD-75375. The non-sheathed filament from the *T. denticola* ATCC 35405 wild-type strain is deposited as PDB: 10PP and EMD-75376. (**Extended Data Table 5**).

## Acknowledgments

We thank Jennifer Aronson for critical reading of the manuscript and Chunyan Wang and Shenping Wu for assisting in cryo-EM data collection. The project was supported by grants R01AI087946, R01AI132818, and R01AI078958 from the National Institute of Allergy and Infectious Diseases (NIAID), R01DE023080 and R01DE034063 from the National Institute of Dental, Craniofacial Research (NIDCR). Cryo-ET data were collected at Yale CryoEM resource, which is funded in part by the NIH grant 1S10OD023603-01A1. We thank the Yale Center for Research Computing facility (YCRC) for providing computing support and advice. We thank the Proteomics and Metabolomics Facility of Cornell University for providing the mass spectrometry data and NIH SIG grant 1S10 OD017992-01 support for the Orbitrap Fusion mass spectrometer.

## Supplemental Information

**Extended Data Fig. 1.**
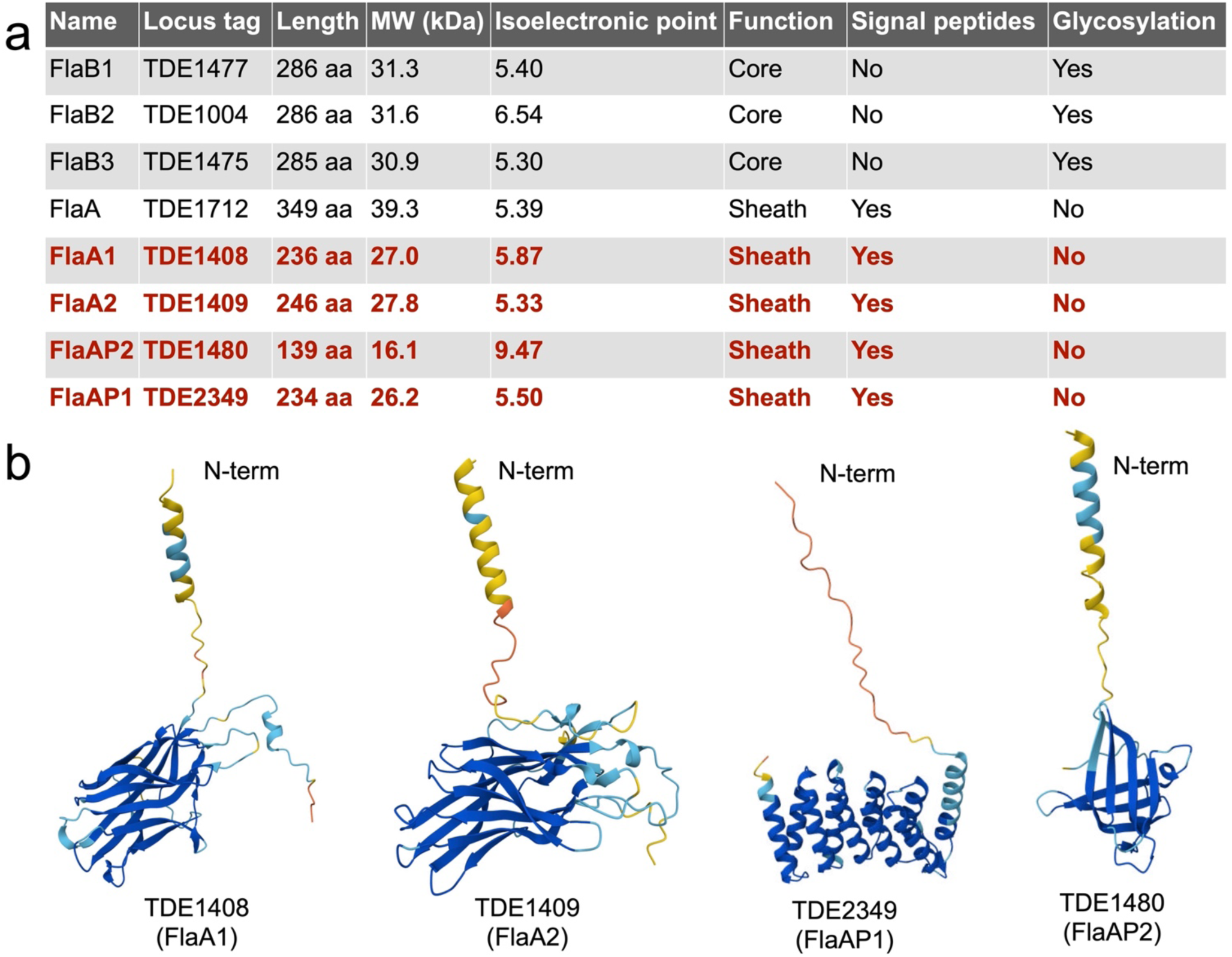
Proteins involved in assembly of the *T. denticola* flagellar filament. **a,** Flagellar filament-associated proteins in *T. denticola*. Proteins colored in red were newly identified in this study. **b,** AlphaFold-predicted structures of FlaA1, FlaA2, FlaAP1, and FlaAP2.

**Extended Data Fig. 2.**
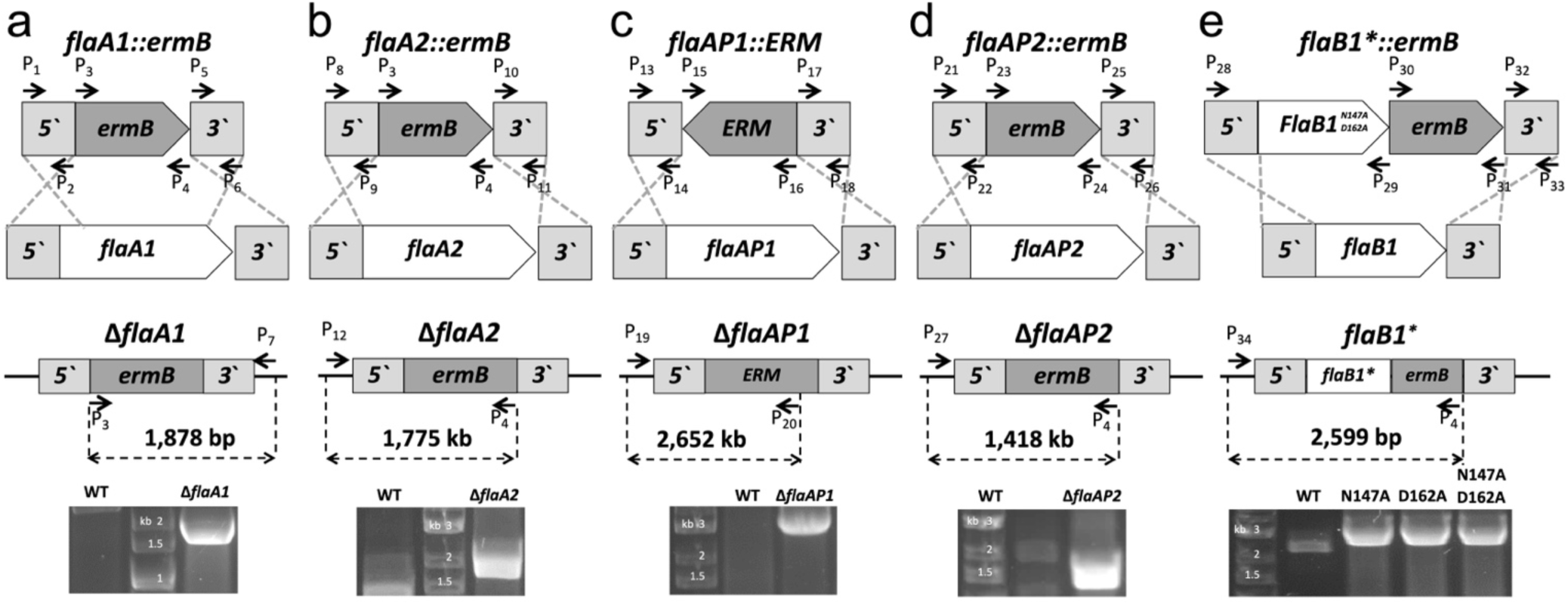
Construction of flagellar mutants. **a-d,** Top panels: Schematic illustrations of the construction of flagellar sheath mutants Δ*TDE1408* (**a**), Δ*TDE1409* (**b**), Δ*TDE2349* (**c**), and Δ*TDE1480* (**d**). Bottom panels: PCR confirmation of the corresponding mutant constructs. **e,** Top panel: Schematic illustration of the construction of *flaB1* site-directed mutants (*flaB1-N147A*, *flaB1-D162A*, and *flaB1-N147A-D162A*). Bottom panel: PCR confirmation of the *flaB1* mutant constructs. The vectors (top panels) were constructed and transformed into *T. denticola*, and the resulting mutants (bottom panels) were verified by PCR. *flaB1* site-directed mutants were further confirmed by DNA sequencing. Arrows indicate the relative positions and orientations of the primers listed in Extended Data Table 1. Numbers (bp) denote the predicted sizes of PCR products generated using the corresponding primers, as illustrated.

**Extended Data Fig. 3.**
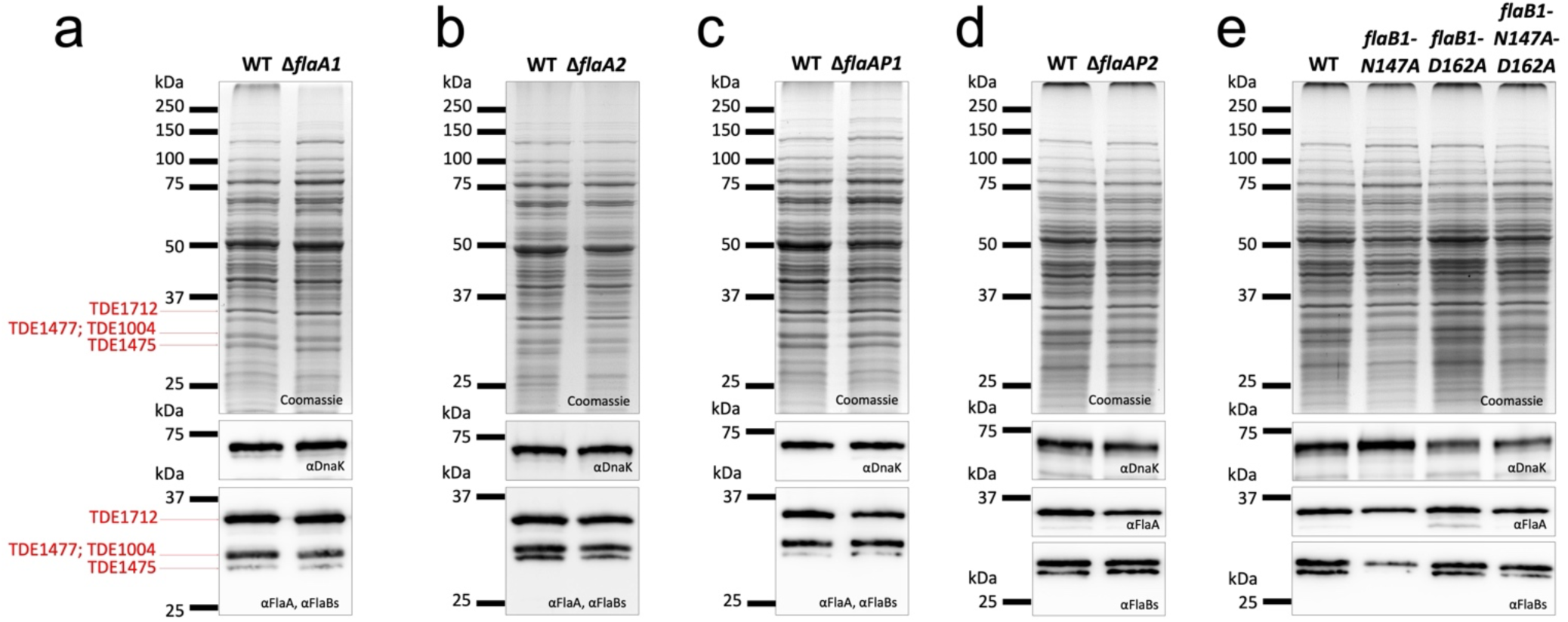
Characterization of flagellar sheath proteins. **a-d,** Whole-cell lysate Coomassie staining (top panel) and immunoblotting (bottom panel) analyses of WT and four flagellar sheath gene deletion mutants, including Δ*flaA1* (**a)**, Δ*flaA2* (**b**), Δ*flaAP1* (**c**), and Δ*flaAP2* (**d**). **e,** Whole-cell lysate Coomassie staining (top panel) and immunoblotting (bottom panel) analyses of WT and *flaB1* site-directed mutants (*flaB1-N147A*, *flaB1-D162A*, and *flaB1-N147A-D162A*). Immunoblots were probed with antibodies against *T. denticola* DnaK (αDnaK), FlaA (αFlaA), and *T. pallidum* FlaB (αFlaB). DnaK was used as a loading control.

**Extended Data Fig. 4.**
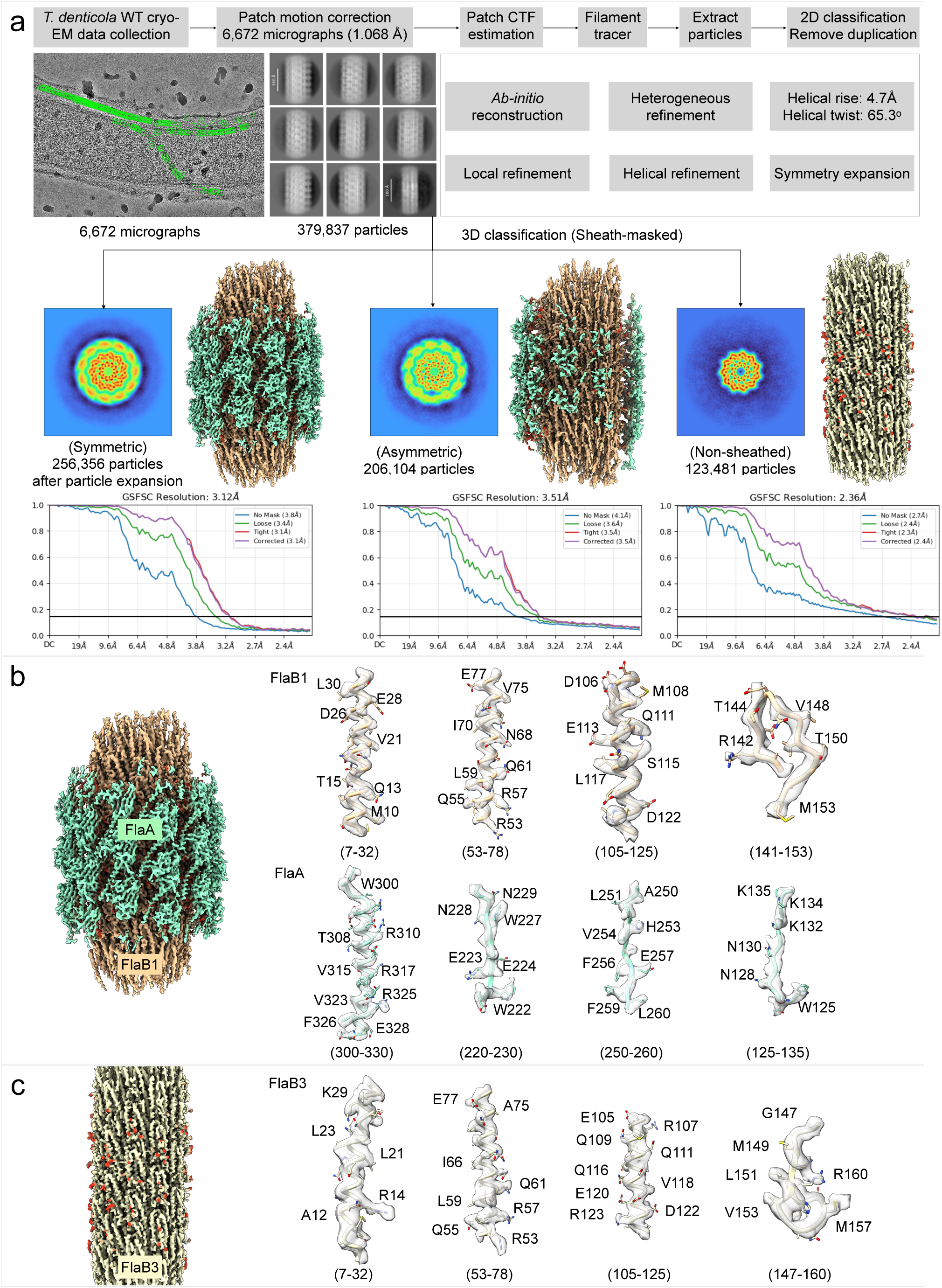
Workflow for *in situ* cryo-EM single-particle analysis of flagellar filaments from wild-type *T. denticola* cells. **a,** A total of 6,672 micrographs were collected from wild-type *T. denticola* cells. Filament segments were automatically picked using the filament tracer tool in cryoSPARC^34^, yielding 42,452 particles after 2D classification. An initial model was generated by *ab initio* reconstruction, followed by helical refinement to obtain an initial symmetric filament structure. The helical rise and twist were determined to be 4.7 Å and 65.3°, respectively. Using the symmetric filament structure as a reference, 3D classification was performed with a sheath mask applied, resulting in three distinct classes corresponding to symmetric, asymmetric, and non-sheathed filaments. Local refinement was subsequently performed to further improve map resolution. **b**, Representative atomic models fitted into cryo-EM density maps. Side chains (sticks) and protein backbones (tubes) are superimposed on the cryo-EM densities rendered as a transparent gray surface. Density maps were visualized in ChimeraX using the volume zone tool with a 2 Å cutoff. Identical model segments of FlaB1 were fitted into their corresponding density maps. **c,** Model-to-map fitting of FlaB3 into the non-sheathed filament structure.

**Extended Data Fig. 5.**
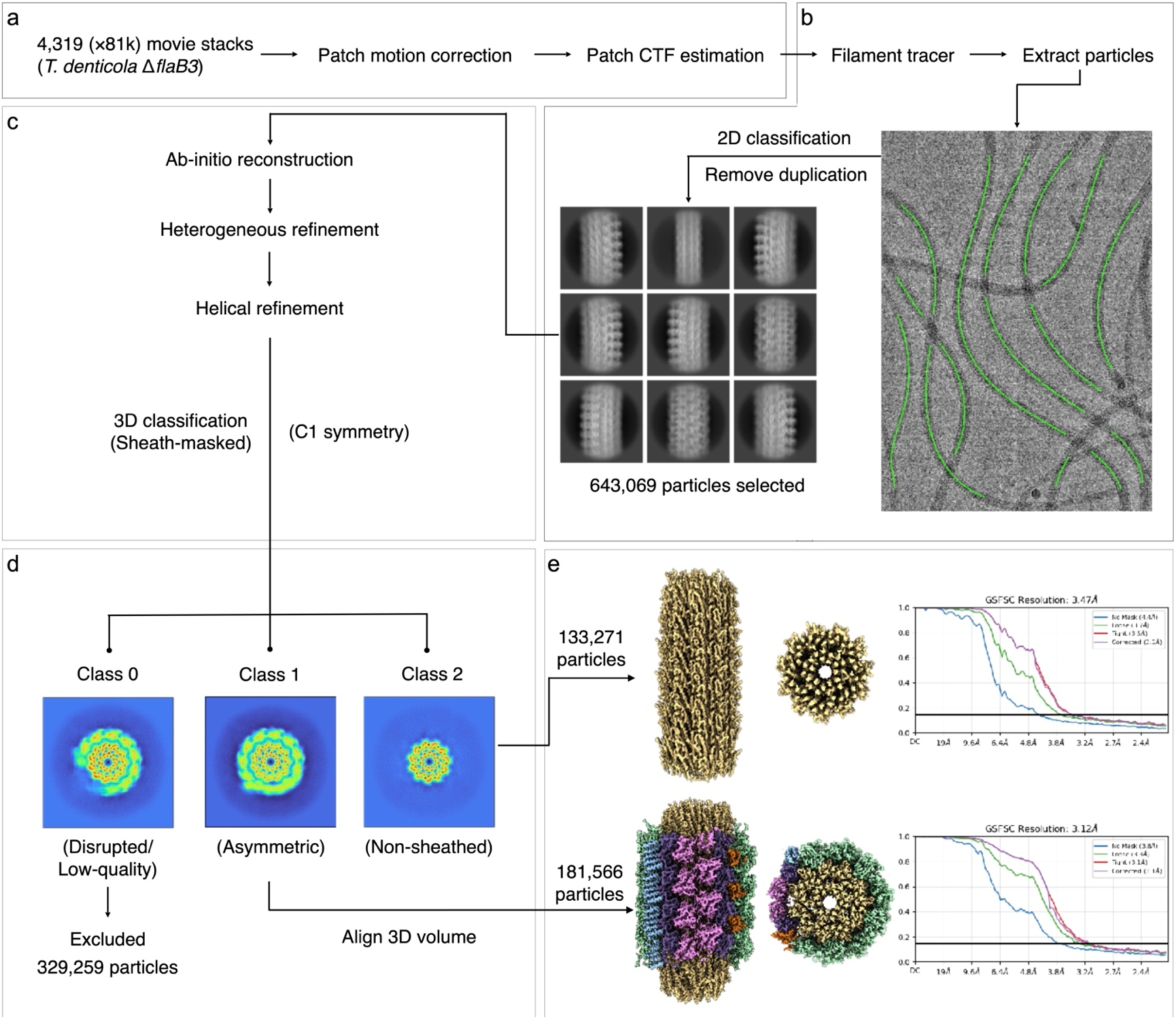
Workflow for *in vitro* single particle cryo-EM of flagellar filaments from the *T. denticola* Δ*flaB3* mutant. A total of 4,319 micrographs were collected from purified flagellar filaments of *T. denticola* Δ*flaB3*. Filament segments were automatically picked using the filament tracer tool in cryoSPARC^34^, yielding 643,069 particles after 2D classification. An initial filament model was generated by homogeneous reconstruction, using a fully symmetric filament volume reconstructed from *in situ* cryo-EM data as the reference. 3D classification was performed using a sheath-focused mask to identify each class. The disrupted or low-quality sheath class was excluded. Non-sheathed filament class was reconstructed by homogeneous refinement. Asymmetric sheath class was further classified using sheath-focused mask by 3D classification and the maps and particles from each class were aligned using the Align 3D Maps in cryoSPARC^34^ until the asymmetric regions were all at a consistent location. All the well-aligned particles were refined by homogeneous refinement. Local refinements were subsequently performed to further improve map resolution.

**Extended Data Fig. 6:**
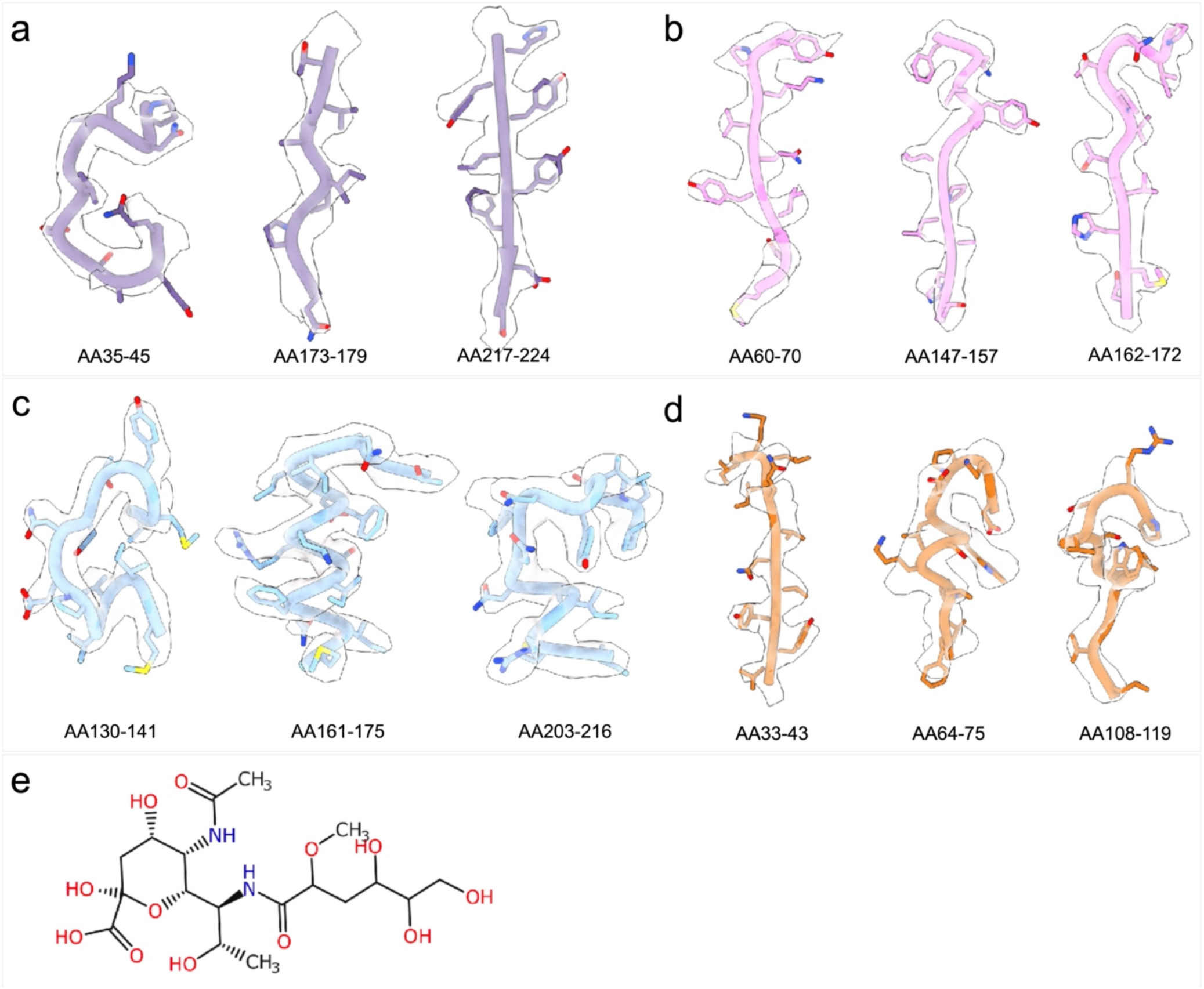
Model-to-map fitting into *in-situ* cryo-EM maps of purified filament from *T. denticola* Δ*flaB3*. **a-d**, Three zoomed-in views of minor sheath proteins. FlaA2 peptide chains (**a**) show the side-chain densities around residues 35-45, 173-179 and 217-224, respectively. FlaA1 peptide chains (**b**) show the side-chain densities around residues 60-70, 147-157 and 162-172, respectively. FlaAP1 peptide chains (**c**) show the side-chain densities around residues 130-141, 161-175 and 203-216, respectively. FlaAP2 peptide chains (**d**) show the side-chain densities around residues 33-43, 64-75 and 108-119, respectively. **e,** The chemical structure of the O-linked glycan modification of FlaB, as previously reported^33^.

**Extended Data Fig. 7:**
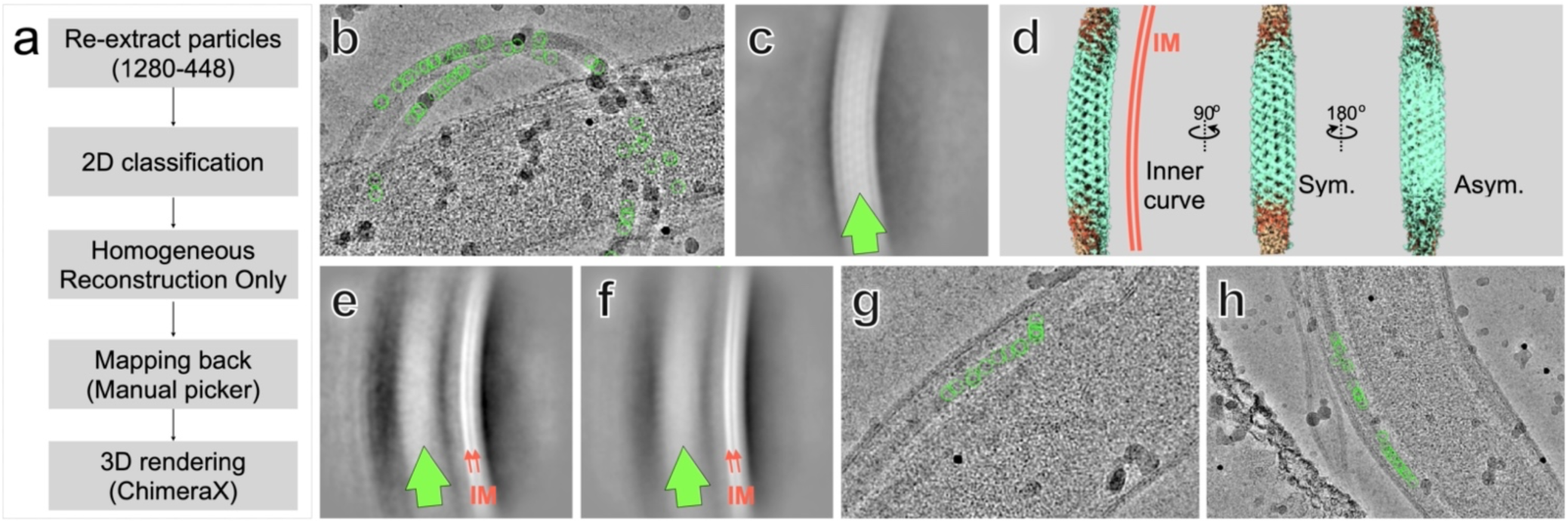
Filament supercoiling in the spirochete *T. denticola*. **a**, Workflow for identification of the asymmetric filament region. **b,** Mapping back of particles exhibiting distinct curvature into raw micrographs. **c**, 2D classification of curvature-selected filament segments. **d,** 3D reconstruction from the same dataset of curvature-selected particles. **e,f,** Representative 2D class averages showing filament segments positioned proximal to the inner membrane, indicating that the asymmetric region is oriented toward the inner membrane of the cell body. **g,h**, Mapping back of particles corresponding to the datasets shown in (**e,f**), confirming their localization near the inner membrane.

**Extended Data Fig. 8.**
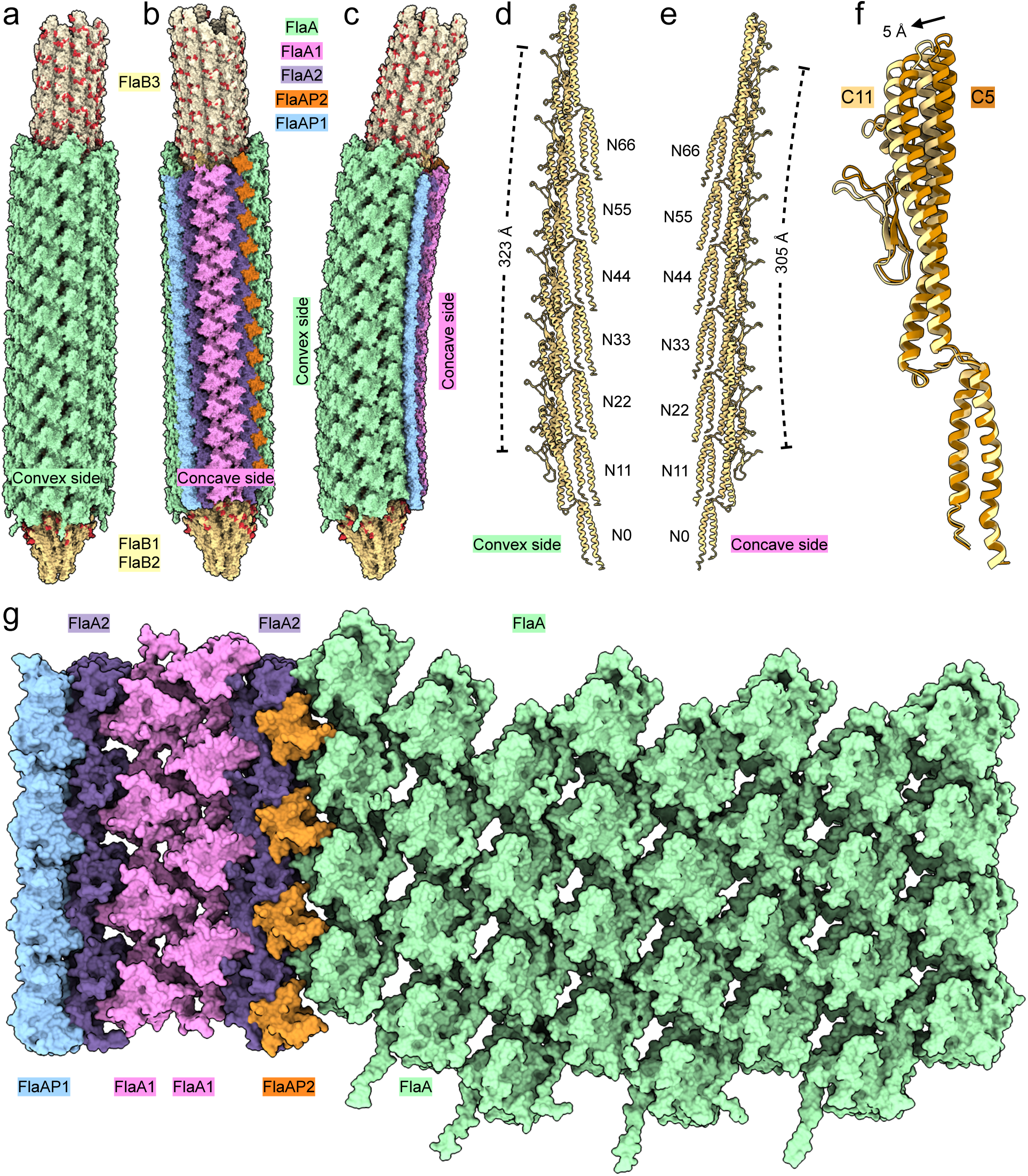
Overall architecture of supercoiled flagellar filament in *T. denticola.* **a-c,** Three different views of the supercoiled flagellar filament. The filament core formed by three core flagellins (FlaB1-FlaB3) is surrounded by the sheath composed of five proteins: FlaA, FlaA1, FlaA2, FlaAP1, and FlaAP2. **d,** The protofilament of FlaB1 at the convex side of the filament. The total length for six FlaA molecules is 323Å. **e,** The protofilament of FlaB1 at the concave side of the filament. The total length for six FlaA molecules is 305Å. **f,** Two FlaB1 molecules from the concave or convex sides are aligned with D0. The D1 domain appears to have 5Å shift. **g,** Unrolled model of the asymmetric filament sheath composes of FlaA, FlaA1, FlaA2, FlaAP2, and FlaAP1, with each component shown in a distinct color.

**Extended Data Fig. 9.**
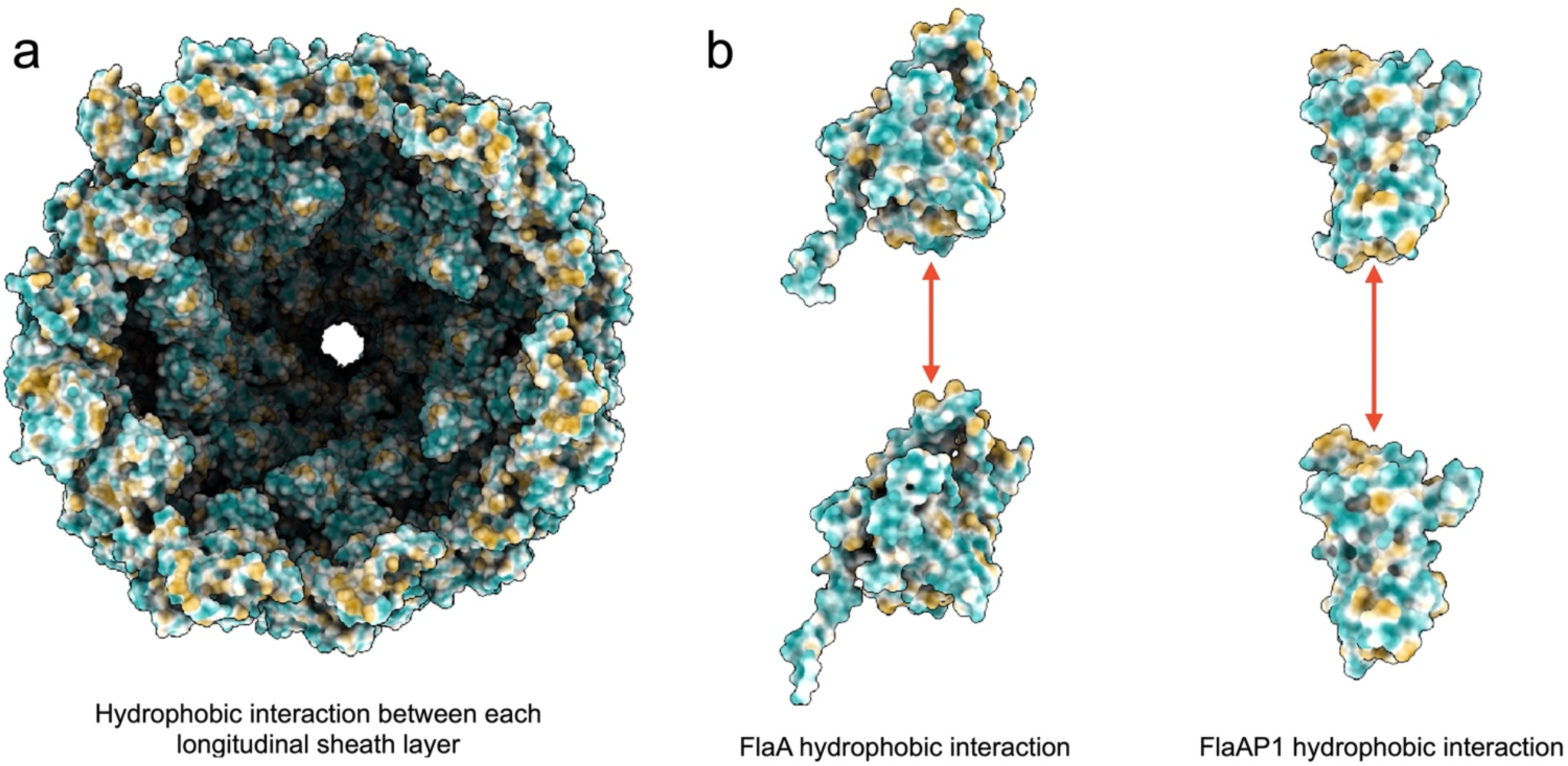
Hydrophobicity analysis of the sheathed filament structure. **a**, Longitudinal cross-sectional view of the sheathed filament highlighting extensive hydrophobic regions at the interlayer interface, which form hydrophobic contacts that contribute to filament structural integrity. **b**, Representative hydrophobic interactions within FlaA longitudinal dimers and FlaAP1 longitudinal dimers.

**Extended Data Fig. 10.**
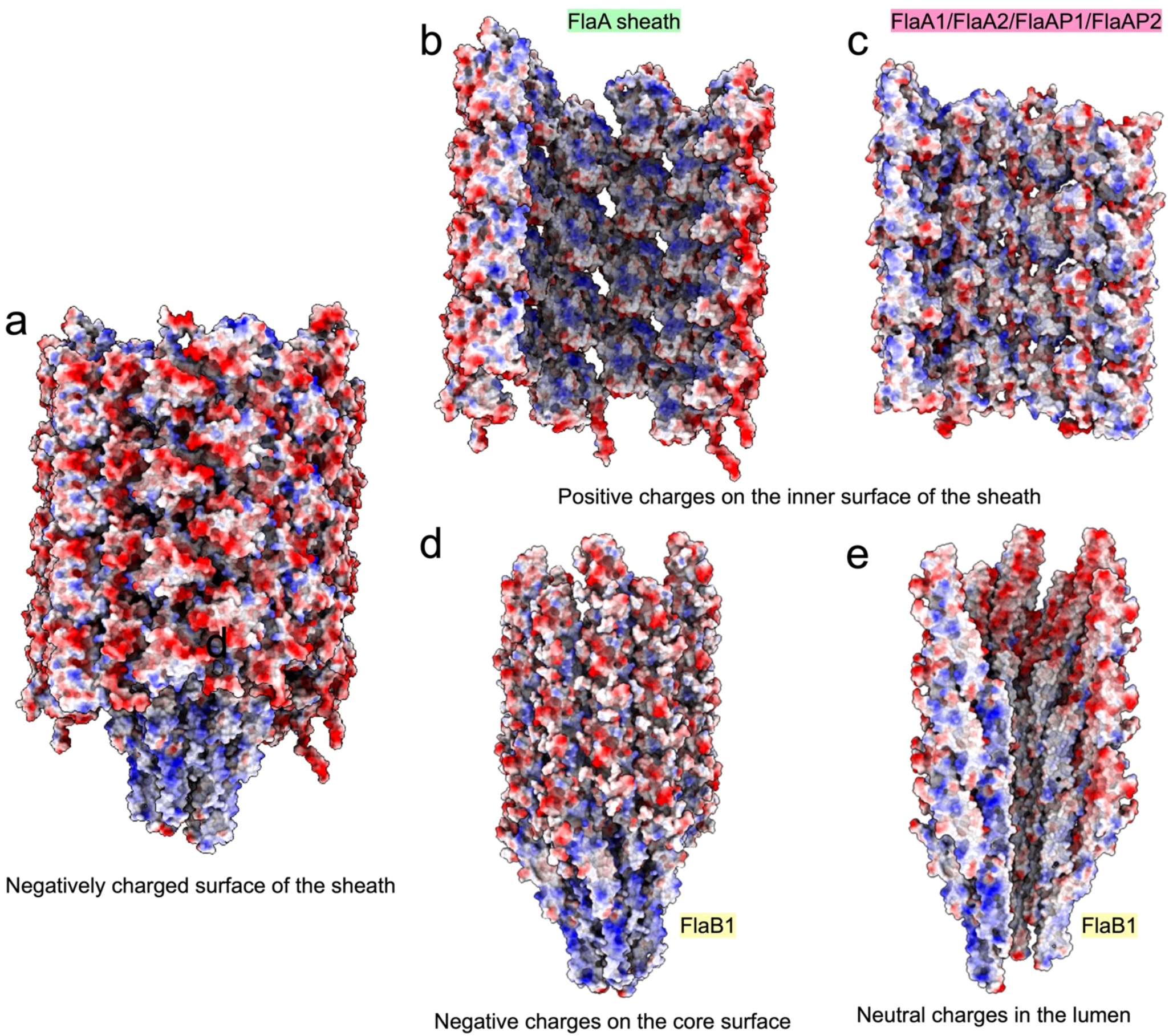
Electrostatic potential analysis of the sheathed filament structure. **a**, Lateral view of the electrostatic potential surface showing that the outer surface of the sheath is predominantly negatively charged, which may generate repulsive interactions with the negatively charged inner membrane and thereby enhance flexibility of flagellar motion. **b,c**, Electrostatic potential mapped onto the sheath surface, revealing a positively charged inner surface in the FlaA region (**b**) and the FlaA1/FlaA2 region (**c**). **d,e**, Electrostatic potential analysis of the glycosylated filament core. A lateral view shows that flexible surface loop regions of the core are predominantly negatively charged (**d**), which may promote attractive interactions with the positively charged inner surface of the sheath, stabilizing core-sheath association and enabling coordinated motion during flagellar rotation. Cross-sectional views show that the lumen of the filament core is relatively electrostatically neutral (**e**), which may facilitate outward secretion of FlaB proteins through the central channel without electrostatic interference with protein conformation or secretion kinetics.

**Extended Data Table 1.**
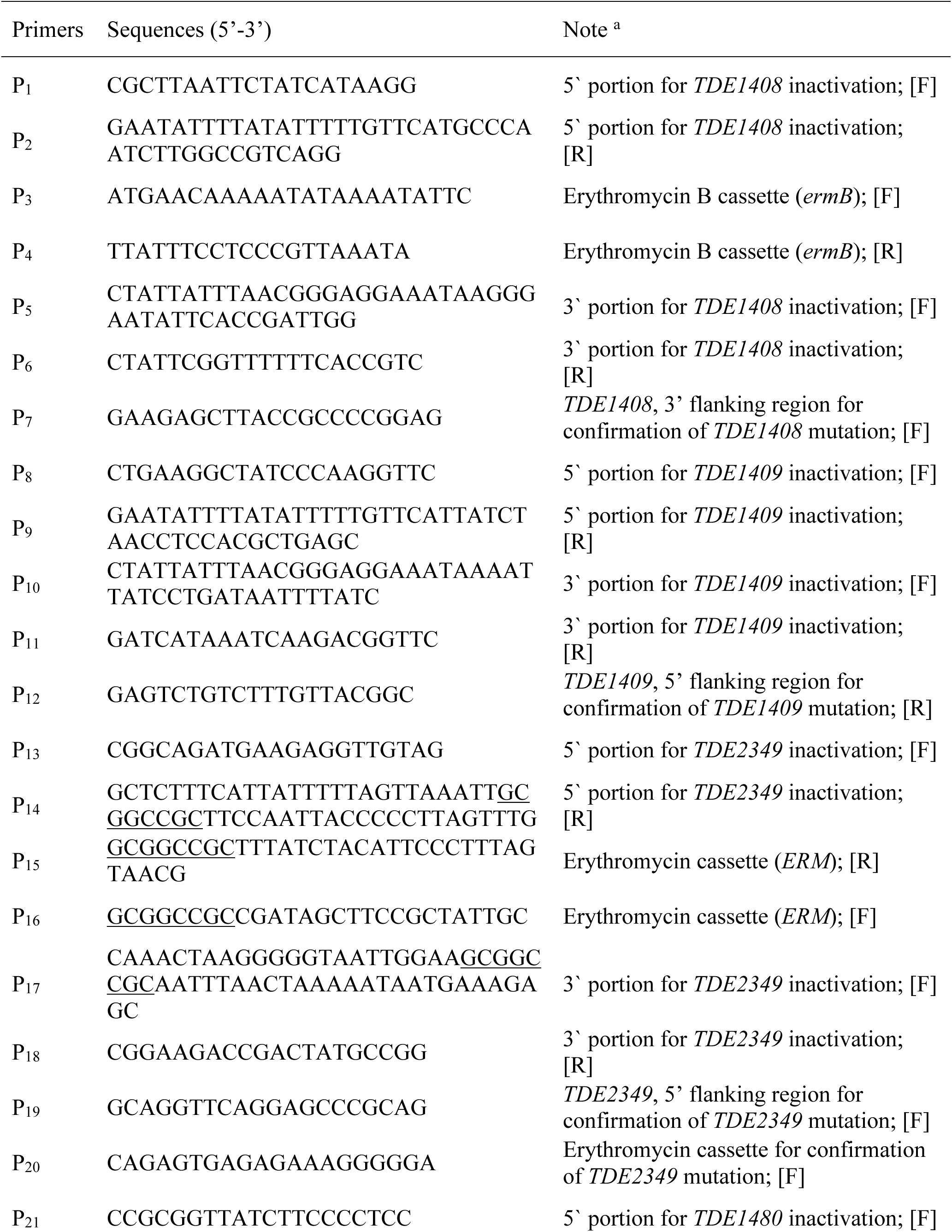
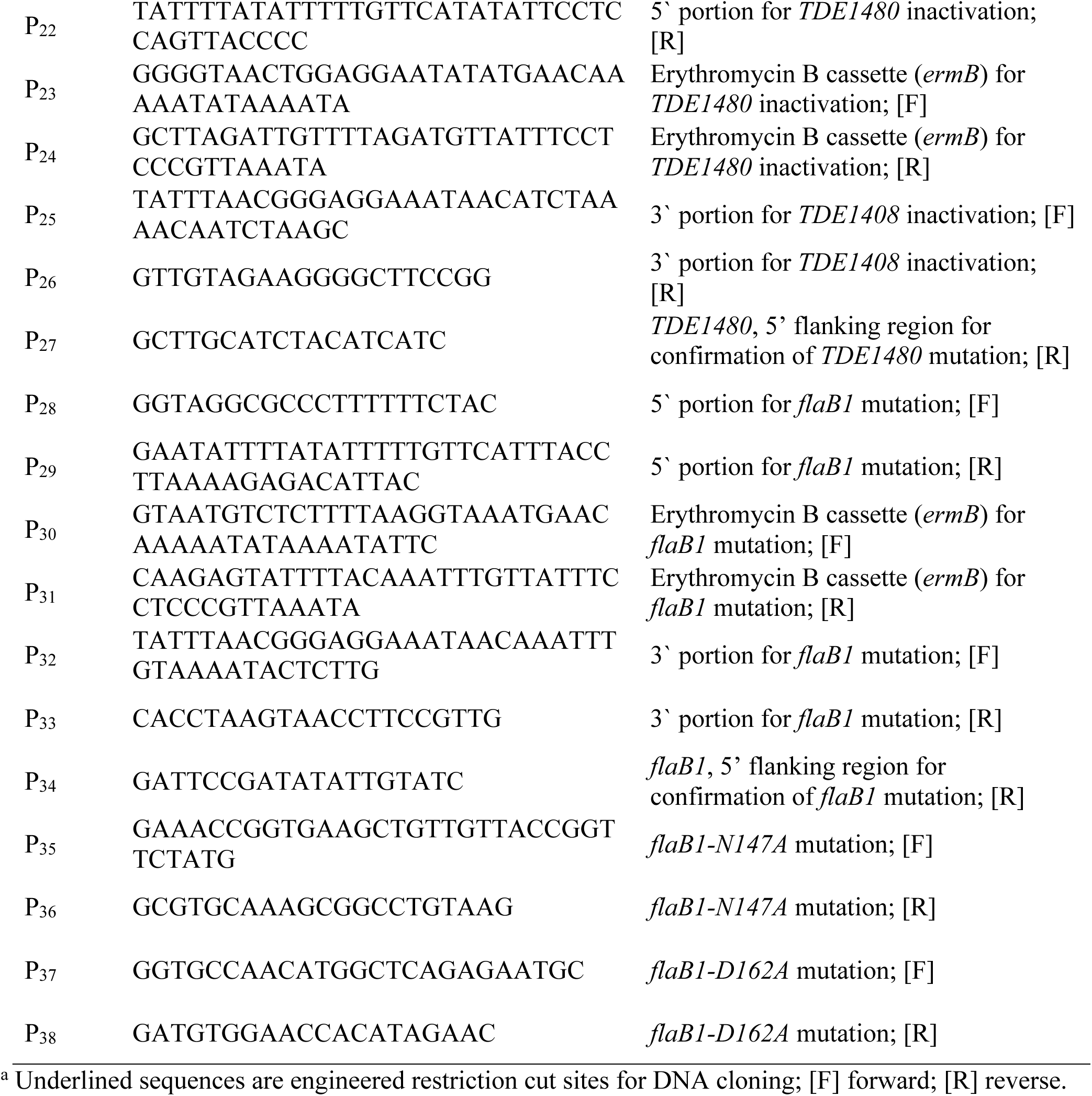
Oligonucleotide primers used in this study.

**Extended Data Table 2.** Stoichiometry of flagellar filament proteins in the flagellar outer sheath and *flaB1* site-directed mutants.

|  | FlaA | FlaBs |
| --- | --- | --- |
| WT | 1 | 1 |
| *ΔflaA1* | 0.946236559 | 0.967741935 |
| WT | 1 | 1 |
| *ΔflaA2* | 0.924731183 | 0.827956989 |
| WT | 1 | 1 |
| *ΔflaAP1* | 0.708737864 | 1.126213592 |
| WT | 1 | 1 |
| *ΔflaAP2* | 0.781609195 | 1.252873563 |
| WT | 1 | 1 |
| *flaB1-N147A* | 0.561983471 | 0.272727273 |
| *flaB1-D162A* | 1.1625 | 1.1375 |
| *flaB1-N147A-D162A* | 0.987179487 | 0.923076923 |

**Extended Data Table 3.**
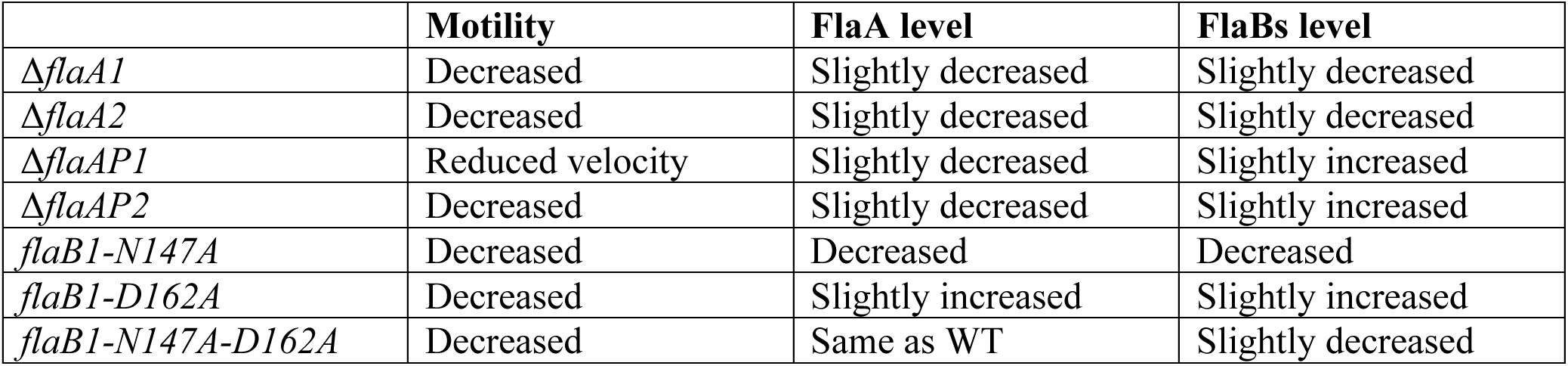
The phenotypes of the flagellar outer sheath and FlaB1 site-directed mutants.

**Extended Data Table 4.** Cryo-ET data of *T. denticola* mutants.

| | $\Delta flaA$ | $\Delta flaA1$ | $\Delta flaA2$ | $\Delta flaAP1$ | $\Delta flaAP2$ | *flaB1-N147A* | *flaB1-D162A* |
| --- | --- | --- | --- | --- | --- | --- | --- |
| Number of tomograms | 7 | 7 | 10 | 11 | 11 | 13 | 12 |
| Magnification | 17,500 | 17,500 | 17,500 | 17,500 | 17,500 | 17,500 | 17,500 |
| Pixel size (Å/pixel) | 1.173 | 1.173 | 1.173 | 1.173 | 1.173 | 1.173 | 1.173 |
| Defocus (µm) | ~4.8 | ~4.8 | ~4.8 | ~4.8 | ~4.8 | ~4.8 | ~4.8 |

**Extended Data Table 5.** Cryo-EM Structure Validation Summary.

| Bacterial strains | $\Delta flaB3$ | | $\Delta flaB3$ | | WT | |
| --- | --- | --- | --- | --- | --- | --- |
| Sheath | Sheathed |  | Non-sheathed |  | Non-sheathed |  |
| Core flagellin | FlaB1 |  | FlaB1 |  | FlaB3 |  |
| Sheath flagellin | FlaA/FlaA1/FlaA2/FlaAP1/FlaAP2 |  |  |  |  |  |
| Refinement | Local |  | Local |  | Local |  |
| PDB | 10PL |  | 10PM |  | 10PP |  |
| EMDB | EMD-75374 |  | EMD-75375 |  | EMD-75376 |  |
| Resolution estimated |  |  |  |  |  |  |
| Metric | Masked (Å) | Unmasked (Å) | Masked (Å) | Unmasked (Å) | Masked (Å) | Unmasked (Å) |
| Using map alone (d99) | 2.53 | 2.57 | 2.89 | 2.93 | 2.67 | 2.70 |
| Overall Biso | 60.00 | 55.00 | 75.00 | 75.00 | 70.00 | 65.00 |
| d_model | 2.20 | 3.90 | 2.90 | 2.90 | 3.20 | 3.30 |
| d_model (B factors = 0) | 4.00 | 4.00 | 4.30 | 4.30 | 4.10 | 4.00 |
| FSC (model) = 0 | 1.74 | 1.85 | 1.90 | 1.94 | 1.84 | 1.91 |
| FSC (model) = 0.143 | 2.21 | 3.91 | 2.91 | 3.84 | 3.19 | 3.50 |
| FSC (model) = 0.5 | 4.11 | 112.49 | 28.95 | 4.70 | 25.21 | 5.36 |
| Mask smoothing radius (Å) | 6.24 |  | 6.94 |  | 4.72 |  |
| Model Validation Statistics |  |  |  |  |  |  |
| Chains | 25 |  | 11 |  | 11 |  |
| Atoms (Hydrogens) | 53988 |  | 24002 |  | 23760 |  |
| Residues | 6889 |  | 3146 |  | 3135 |  |
| RMSD bond lengths (Å) | 0.008 |  | 0.017 |  | 0.018 |  |
| RMSD bond angles (°) | 1.320 |  | 1.953 |  | 2.197 |  |
| MolProbity score | 1.96 |  | 2.36 |  | 3.06 |  |
| Clash score | 6.82 |  | 19.44 |  | 22.31 |  |
| Ramachandran favored (%) | 94.50 |  | 96.80 |  | 88.37 |  |
| Ramachandran allowed (%) | 5.16 |  | 2.53 |  | 8.99 |  |
| Ramachandran outliers (%) | 0.34 |  | 0.67 |  | 2.63 |  |
| Rama-Z (whole / helix / sheet / loop) | -3.03 / -2.30 / -0.81 / -2.05 |  | 0.62 / 1.36 / - / -2.00 |  | -1.05 / 0.15 / - / -2.49 |  |
| Rotamer outliers (%) | 1.9 |  | 3.16 |  | 6.69 |  |
| CaBLAM outliers (%) | 3.20 |  | 1.19 |  | 3.69 |  |
| Occupancy mean | 1.00 |  | 1.00 |  | 1.00 |  |
| B-factor (mean / min / max) | 88.66 / 28.65 / 254.87 |  | 102.62 / 56.36 / 176.82 |  | 66.79 / 19.72 / 152.67 |  |
| CC (mask / box / peaks / volume) | 0.70 / 0.31 / -0.06 / 0.70 |  | 0.72 / 0.32 / -0.07 / 0.72 |  | 0.68 / 0.35 / 0.04 / 0.68 |  |

